# Cdc48 Cofactor Shp1 Regulates Signal-Induced SCF^Met30^ Disassembly

**DOI:** 10.1101/2019.12.13.876029

**Authors:** Linda Lauinger, Karin Flick, James L. Yen, Radhika Mathur, Peter Kaiser

**Affiliations:** Department of Biological Chemistry, School of Medicine, 240D Medical Science I University of California, Irvine, CA 92697-1700, USA; NeoGenomics Laboratories; DiaSorin Molecular LLC

## Abstract

Organisms can adapt to a broad spectrum of sudden and dramatic changes in their environment. These abrupt changes are often perceived as stress and trigger responses that facilitate survival and eventual adaptation. The ubiquitin proteasome system (UPS) is involved in most cellular processes. Unsurprisingly, components of the UPS also play crucial roles during various stress response programs. The budding yeast SCF^Met30^ complex is an essential Cullin-RING ubiquitin ligase that connects metabolic and heavy metal stress to cell cycle regulation. Cadmium exposure results in the active dissociation of the F-box protein Met30 from the core ligase leading to SCF^Met30^ inactivation. Consequently, SCF^Met30^ substrate ubiquitylation is blocked and triggers a downstream cascade to activate a specific transcriptional stress response program. Signal-induced dissociation is initiated by autoubiquitylation of Met30 and serves as a recruitment signal for the AAA-ATPase Cdc48/p97, which actively disassembles the complex. Here we show that the UBX cofactor Shp1/p47 is an additional key element for SCF^Met30^ disassembly during heavy metal stress. Although the cofactor can directly interact with the ATPase, Cdc48 and Shp1 are recruited independently to SCF^Met30^ during cadmium stress. An intact UBX domain is crucial for effective SCF^Met30^ disassembly, and a concentration threshold of Shp1 recruited to SCF^Met30^ needs to be exceeded to initiate Met30 dissociation. The latter is likely related to Shp1-mediated control of Cdc48 ATPase activity. This study identifies Shp1 as the crucial Cdc48 cofactor for signal-induced, selective disassembly of a multi-subunit protein complex to modulate activity.

**Significance Statement:** Ubiquitylation affects many important cellular processes, and has been linked to a number of human diseases. It has become a synonym for protein degradation, but ubiquitylation also has important non-proteolytic signaling functions. Understanding the molecular concepts that govern ubiquitin signaling is of great importance for development of diagnostics and therapeutics. The cadmium-induced inactivation of the SCF^Met30^ ubiquitin ligase via the disassembly of the multi-subunit ligase complex, illustrates an example for non-proteolytic signaling pathways. Dissociation is triggered by autoubiquitylation of the F-box protein Met30, which is the recruiting signal for the highly conserved AAA-ATPase Cdc48/p97. Here we show that the UBX cofactor Shp1/p47 is important for this ubiquitin-dependent, active remodeling of a multi-protein complex in response to a specific environmental signal.

## Introduction

The maintenance of protein homeostasis is crucial for eukaryotic cells. The post-translational modification of a protein by ubiquitin conjugation was first described as the major non-lysosomal mechanism by which proteins are targeted for degradation (1). However, destruction of proteins via the proteasomal pathway is only one of many outcomes of ubiquitylation. The fate of a protein is decided depending on how many ubiquitin molecules are covalently attached, and in which fashion ubiquitin chains are formed. The outcomes can be versatile, and, for example, influence abundance, activity, or localization of proteins (2–5). The process of ubiquitin conjugation requires the coordinated reaction of the E1-E2-E3 enzyme cascade. E3 ligases mediate the final step of substrate specific conjugation (6–8). E3s are the most diverse components in the ubiquitylation machinery and are divided in several classes. The largest group is represented by the multi-subunit cullin-RING ligases (CRLs), which include the well-studied subfamily of SCF (Skp1-cullin-F-box) ligases (9, 10). SCFs are composed of four principal components: The RING finger protein Rbx1, the scaffold Cul1, and the linker Skp1 that forms an association platform with different substrate-specific F-box proteins (11, 12).

The punctual degradation of several SCF substrates is essential to ensure normal cell growth. Hence, alterations in SCF component expression or function can often be linked to cancer and other diseases (13, 14). This highlights the importance to better understand SCF ligase regulation and enable their therapeutic targeting. It has long been thought that ubiquitylation by SCF ligases is solely regulated at the level of substrate binding (11, 15). However, an additional mode of CRL regulation was recently discovered. Specific SCF ligases can be inhibited by signal-induced dissociation of the F-box subunit from the core ligase (16, 17). The best studied example is SCF^Met30^, which controls ubiquitylation of a number of different substrates (16, 18–20). The transcriptional activator Met4 and the cell-cycle inhibitor Met32 are the most critical ones. Together they orchestrate induction of cell cycle arrest and activation of a specific transcriptional response during nutritional or heavy metal stress. This stress response protects cellular integrity, restores normal levels of sulfur containing metabolites, and activates a defense system for protection against heavy metal stress (21–26). Under normal growth conditions SCF^Met30^ mediates Met4 and Met32 ubiquitylation. Even though both are modified with the classical canonical destruction signal, a lysine-48 linked ubiquitin chain, only Met32 is degraded via the proteasomal pathway. In contrast Met4 is kept in an inactive state by the attached ubiquitin chain (27, 28). Both, nutrient and heavy metal stress block SCF^Met30^ dependent ubiquitylation of Met4 and Met32, however through profoundly distinctive mechanisms. Nutrient stress prevents the interaction between the SCF^Met30^ ligase and its substrate following the canonical mode of regulation, but cadmium stress leads to active dissociation of the F-box subunit Met30 from the core ligase (21, 22). Remarkably, dissociation is initiated by autoubiquitylation of the F-box protein Met30, which serves as the recruiting signal for the conserved AAA-ATPase Cdc48/p97 (16). Importantly, ubiquitin-dependent recruitment of Cdc48 and dissociation of Met30 from the core ligase is independent of proteasome activity (16).

Cdc48 is involved in many diverse cellular pathways including extraction of unfolded proteins from the ER (29, 30), discharge of membrane bound transcription factors (31, 32), and chromatin-associated protein extraction (33–38). However, Cdc48 itself rarely displays substrate specificity. An ensemble of regulatory cofactors tightly control functions of the ATPase by recruiting it to different cellular pathways (39–41). Shp1 (**S**uppressor of **H**igh copy protein **P**hosphatase 1 or p47 in mammals) is part of the UBA-UBX family of Cdc48 cofactors (42). The UBA domain binds ubiquitin, with a preference for multi-ubiquitin chains (43, 44), and the UBX domain is structurally similar to ubiquitin and directly interacts with Cdc48 by mimicking a mono-ubiquitylated substrate (45–48). An additional domain, the SEP (**S**hp1/**E**yeless/**P**47) domain, is involved in trimerization of the cofactor (49, 50). N-terminal of the UBX domain, a short hydrophobic sequence in Shp1 represents an additional Cdc48 binding motif (BS1), which enables a bipartite mode of Shp1 binding to Cdc48 (41, 46, 51, 52).

In yeast, Shp1 is involved in Cdc48-dependent protein degradation via the UPS (45). The Cdc48^Shp1^ complex also interacts with the ubiquitin-fold autophagy protein Atg8, and is a component of autophagosome biogenesis (53). The best studied function of Shp1 relates to activation of protein phosphatase 1 (Glc7), which is important for cell cycle progression (54, 55). Hence, deletion of Shp1 leads to a severe growth phenotype and cells are prone to genomic instability (45, 55, 56). Here we show that Shp1 is the crucial Cdc48 cofactor for SCF^Met30^ disassembly in response to heavy metal stress. We identified the very C-terminal 22 amino acids to be important for disassembly. Surprisingly, Cdc48 and Shp1 get independently recruited to SCF^Met30^ during cadmium stress, and recruitment depends on SCF^Met30^ autoubiquitylation. Furthermore, initiation of SCF^Met30^ disassembly is triggered by a Shp1 threshold level likely necessary for stimulating ATP hydrolysis by Cdc48 (57).

## Results

### Shp1 is involved in the Cellular Response during Cadmium Stress

The distinct functions of Cdc48/p97 in diverse cellular processes are tightly controlled by a large number of cofactors (40). For mechanical reasons Cdc48 requires at least two anchor points to generate physical force and make Met30 dissociation possible. We reasoned that deletion of a cofactor necessary for SCF^Met30^ disassembly would result in cadmium sensitivity, because defense pathways would not be activated. Deletion of *SHP1* rendered yeast cells cadmium sensitive (16, 45), and we therefore considered Shp1 as a potential cofactor for SCF^Met30^ disassembly. This hypothesis was supported by immunoprecipitation experiments of Met30 that showed Cdc48 and Shp1 are recruited to SCF^Met30^ in response to cadmium exposure (Fig 1A). We have previously demonstrated that Met30 dissociates rapidly from Skp1 concomitant with Cdc48 recruitment to SCF^Met30^ when cells are exposed to cadmium (16) and Fig1A). In *shp1Δ* deletion mutants (K.O.) Met30 dissociation kinetics were severely delayed. Additionally, the amount of Cdc48 in complex with Met30 in the absence of Shp1 was significantly increased even under unstressed growth conditions, and accumulated to even higher levels during cadmium stress (Fig1A, Suppl. Fig 1A). Increased Cdc48 binding suggests that the dissociation complex intermediate is trapped in *shp1Δ* mutants, and importantly, that Shp1 is not required for Cdc48 recruitment, but to trigger Cdc48-catalyzed Met30 dissociation (Fig1A, Suppl. Fig 1A).

**Figure 1.**
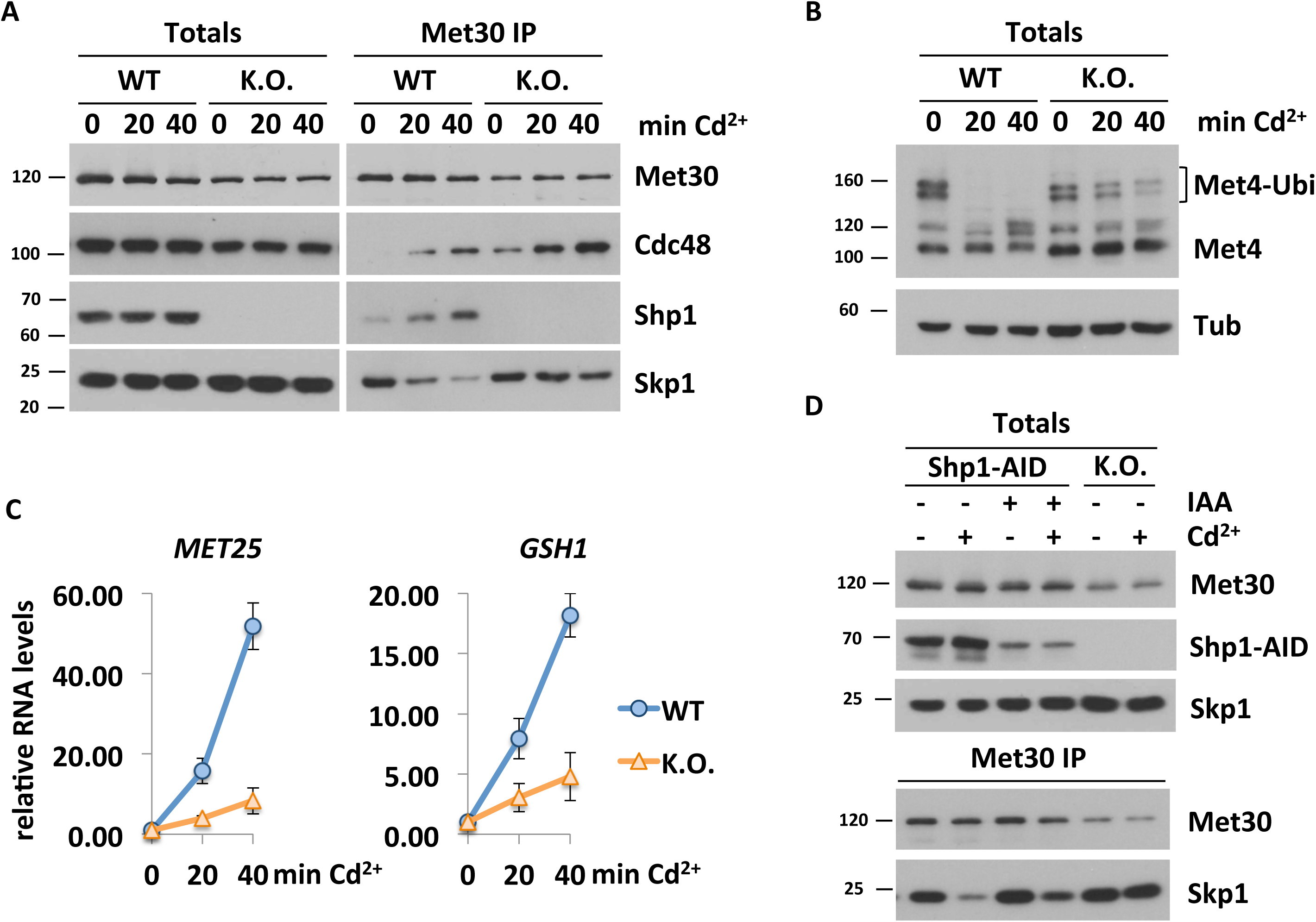
Shp1 is involved in the Cellular Response during Cadmium Stress. A) Shp1 is recruited to SCF^Met30^ during Cadmium Stress. Dissociation kinetics of Met30 from the SCF core ligase are decreased in *shp1Δ* deletion mutants (K.O.) Strains expressing endogenous ^12xMyc^Met30, Cdc48^RGS6H^, Skp1, and Shp1^3xHA^ in WT or *shp1Δ* cells were cultured at 30°C in YEPD medium and treated with 100 µM CdCl_2_ and samples were harvested at indicated time points. ^12xMyc^Met30 was immunoprecipitated and co-precipitated proteins were analyzed by Western blot. B) Shp1 is necessary for Met4 activation during Heavy Metal Stress. Whole cells lysates (Totals) of samples shown in Figure 1A were analyzed by Western blot using a Met4 antibody to follow the ubiquitylation and phosphorylation status of Met4. Tubulin was used as a loading control. C) Cadmium induced gene expression is altered in the absence of Shp1. RNA was extracted from samples shown in Figure 1A. Expression of Met4 target genes *MET25* and *GSH1* was analyzed by RT-qPCR and normalized to 18S rRNA levels (n=3), data are represented as mean ±SD. D) Temporal down-regulation of Shp1 shows its importance during Cadmium Stress. Strains expressing endogenous ^12xMyc^Met30, Shp1^3xHA-AID^ and the F-Box protein ^2xFLAG^OsTir under the constitutive ADH promoter were cultured at 30°C in YEPD medium in the absence and presence of 500 µM auxin for 4 hours to deplete endogenous Shp1. Cells were treated with 100 µM Cadmium and samples were harvested after 20 min of heavy metal exposure. ^12xMyc^Met30 was immunoprecipitated and co-precipitated proteins were analyzed by Western blot. Results shown in A, B and D are representative blots from three independent experiments.

Under normal growth conditions SCF^Met30^ facilitates the ubiquitylation of Met4 and thereby represses the cadmium related stress response (21, 22). Cadmium-triggered dissociation of Met30 leads to SCF^Met30^ inhibition and thereby blocks Met4 ubiquitylation, leading to cell cycle arrest and induction of the transcription of glutathione and sulfur amino acid biosynthesis regulating genes (25). Consistent with Shp1 requirement for SCF^Met30^ disassembly, Met4 remained ubiquitylated during cadmium stress in *shp1Δ* mutants (Fig 1B). Furthermore, the lack of Met4 activation in the absence of Shp1 is reflected in blunted cadmium-induced Met4 target gene activation (Fig 1C). These results and cadmium hypersensitivity of *shp1Δ* cells (16, 45) suggest, that Shp1 is critical for the Met4-mediated cadmium stress program. However, Shp1 is essential for normal mitotic progression and *shp1Δ* mutants grow very slowly ((55), Suppl Fig 1B) or are inviable in certain genetic backgrounds (45). Thus, as with all mutations that severely compromise cell proliferation, we were concerned that *shp1Δ* deletion strains acquired compensatory mutations that could lead to misinterpretation of results (58). Therefore, we decided to use the *Auxin Inducible Degron* (AID) system (59, 60) to acutely down-regulate Shp1 protein levels. Auxin (IAA) mediates the interaction of the F-box protein Tir1 and an AID-tagged protein leading to the ubiquitylation and proteasomal degradation of the AID-tagged substrate. Shp1 was tagged with a 3xHA-AID-fragment and co-expressed with OsTir in the presence of IAA. Within 30 min of IAA supplementation Shp1-AID levels were significantly decreased (Suppl. Figure 1 C). Even though Shp1 levels were severely down-regulated for several hours, no significant growth defect was observed (Suppl. Fig 1D), indicating, that the remaining Shp1 amount was enough to fulfill its function in mitotic progression, or that genomic instability associated with *shp1Δ* mutants leads to accumulation of growth inhibiting mutations over time. We compared cadmium-induced Met30 dissociation between *shp1Δ* deletion mutants and auxin-mediated knock down strains. When Shp1 levels were low (< 10% of endogenous amount), cadmium-induced Met30 dissociation was compromised. Although, this dissociation phenotype was not as pronounced as in the complete absence of Shp1 (Fig 1D, Suppl. Fig 1E), these results confirm an important role of Shp1 in SCF^Met30^ disassembly during heavy metal stress.

### Mutation of Functional Domains in Shp1

Several conserved functional domains have been characterized in Shp1 and its mammalian orthologue p47 (41). These include the UBA and UBX domains that characterize this family of Cdc48 cofactors. We wanted to evaluate the importance of these domains and the role of Shp1/Cdc48 interaction for SCF**^Met30^** disassembly. The crystal structure of p97 bound to p47, the human orthologs of yeast Cdc48 and Shp1, respectively, has been reported (61). Several regions of p47 contact p97 in this structure. Most notably the S3/S4 loop (residues 342–345 in p47 and 396-398) located in the UBX domain inserts into a hydrophobic pocket formed between two p97 N subdomains. Additional contacts are formed by evolutionary conserved residues at the very C-terminus of p47 (61). Furthermore, we deleted another evolutionary conserved region (residues 304-314 in Shp1) referred to as binding site 1 (BS1) or SHP box, a motif that mediates binding of other proteins to Cdc48 (51, 55, 62, 63). To simplify the nomenclature, we refer to BS1 as Cdc48 interacting domain 1 (CIM1), to the S3/S4 loop as CIM2, and to the last 22 residues of Shp1 as UBX_Ct_ (Fig 2A and Suppl. Fig 2A). The latter two binding elements are embedded in the UBX domain, but mutation of one domain does not interfere with integrity of the other domain. All mutations were generated at the endogenous locus by CRISPR/Cas9 mediated gene editing. The CIM1 mutation was generated by deleting residues 304 to 314, and the CIM2 mutation by changing the important FPI sequence to GAG. As previously reported (55) mutation of either the CIM1, CIM2, or UBX_Ct_ severely reduced steady state interaction of endogenous proteins with Cdc48, while deletion of the UBA domain had no effect (Fig 2D and Suppl. Fig 2E). Mutant Shp1 proteins were expressed at similar levels, and in contrast to *shp1Δ* deletion mutants, none of the mutant strains showed substantial growth defects (Fig 2B&C and Suppl. Fig 2C&D). We noticed significantly reduced Met30 levels in *shp1Δ* mutants, which was not observed in any of the domain mutations (Fig. 2C). Met30 protein stability was not affected, but *shp1Δ* mutants showed reduced *MET30* RNA expression (Suppl. Fig 2F & G). The reason for the expression effect is unknown.

**Figure 2.**
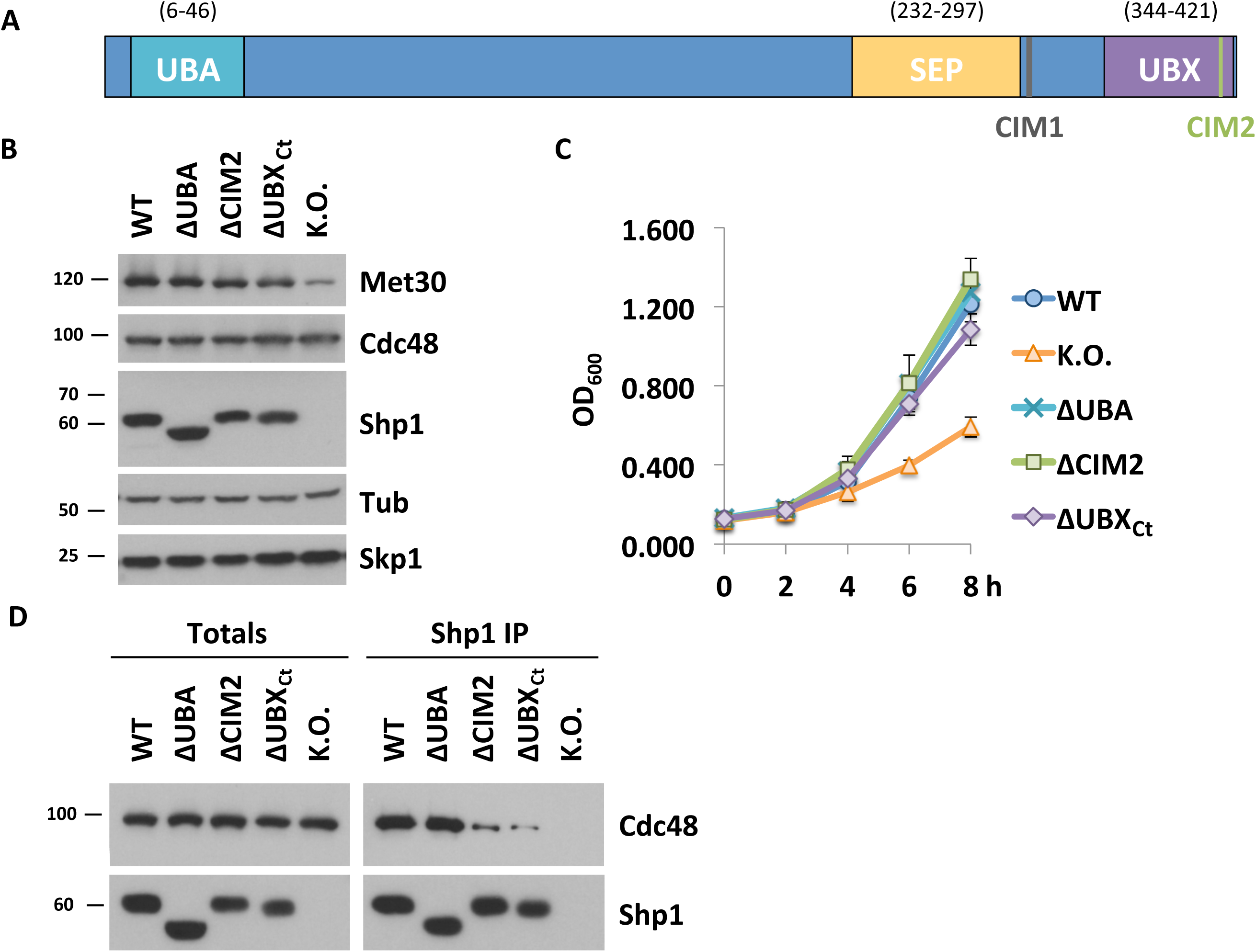
Characterization of Functional Domains in Shp1. A) Schematic of Shp1 and its known functional domains and motifs. UBA = ubiquitin-associated domain (6-46), SEP = Shp1, eyeless and p47 domain (238-313), UBX = ubiquitin regulatory X domain (346-420), CIM 1= Cdc48-interacting motif 1 (LGGFSGQGQRL; 304-314 distal of SEP domain also known as BS1), and CIM2= Cdc48-interacting motif 1 (FPI; 396-398 in the UBX domain). B) Expression levels of Shp1 mutants. Steady-state levels of indicated proteins were compared from strains expressing endogenous ^12xMyc^Met30, Cdc48^RGS6H^, Skp1, and the various Shp1 mutants as indicated (all 3xHA tagged). ΔUBA = deletion of residues 1-50, ΔCIM2 = 396-398 FPI residues were replaced with GAG, ΔUBX_Ct_ = deletion of aa 401-423, or *shp1* gene knocked out (K.O.). Proteins were analyzed by Western blotting with tubulin as a loading control. C) Shp1 mutants do not show a significant growth defect at 30°C. Wild-type cells, *shp1* mutant variants as indicated, and *shp1Δ* (K.O.) strains were grown at 30°C in YEPD medium and samples were taken at indicated time points to measure optic density at 600nm. D) Cdc48 binding is significantly decreased in ΔCIM2 and ΔUBX_Ct_ Shp1 mutants. SHP1^3xHA^ variants were immunoprecipitated and co-precipitation of Cdc48^RGS6H^ was analyzed by Western blotting. The Shp1 K.O. strain was used as a background control. Results shown in B and D are representative blots from three independent experiments.

### The UBX domain of Shp1 is required for SCF^Met30^ Disassembly during Cadmium Stress

We next analyzed the impact of different Shp1 mutants on the response to cadmium stress. We focused these studies on Shp1ΔCIM2, ΔUBA, and ΔUBX_Ct_ mutants, because Shp1 with either ΔCIM1 and ΔCIM2 mutations showed identical behavior in initial experiments (Suppl. Fig 2). As previously shown, the *shp1Δ* deletion mutants were hypersensitive towards cadmium exposure (16, 45) (Fig. 3A). ΔUBX_Ct_ mutants were also cadmium sensitive, albeit only moderately, whereas all other Shp1 domain mutants tolerated cadmium exposure (Fig 3A). Cadmium sensitivity caused by defects in the SCF^Met30^ system is due to a lack of Met4 activation and the resulting blocked induction of a defense program mediated by Met4-dependent gene transcription. Met4 activation can be monitored by appearance of deubiquitylated Met4 and induction of genes such *MET25* and *GSH1* in response to cadmium stress (16, 21, 64). These are sensitive indicators to monitor functionality of the SCF^Met30^ system. Consistent with the cadmium sensitivity assay, *shp1Δ* and *Δubx_Ct_* mutants were unable to fully activate Met4 as demonstrated by maintained ubiquitylated Met4 during cadmium stress (Fig 3B). All other Shp1 variants responded similar to wild-type cells and completely blocked Met4 ubiquitylation (Fig 3B). These findings were further confirmed by analysis of cadmium-induced expression of Met4 target genes. *shp1-ΔUBA* and *shp1-ΔCIM2* showed induction of *MET25* and *GSH1* similar to wild-type Met4. However, *shp1-Δubx_Ct_* mutants failed to efficiently activate gene expression, consistent with persistent Met4 ubiquitylation and cadmium sensitivity of this mutant (Fig 3C).

**Figure 3.**
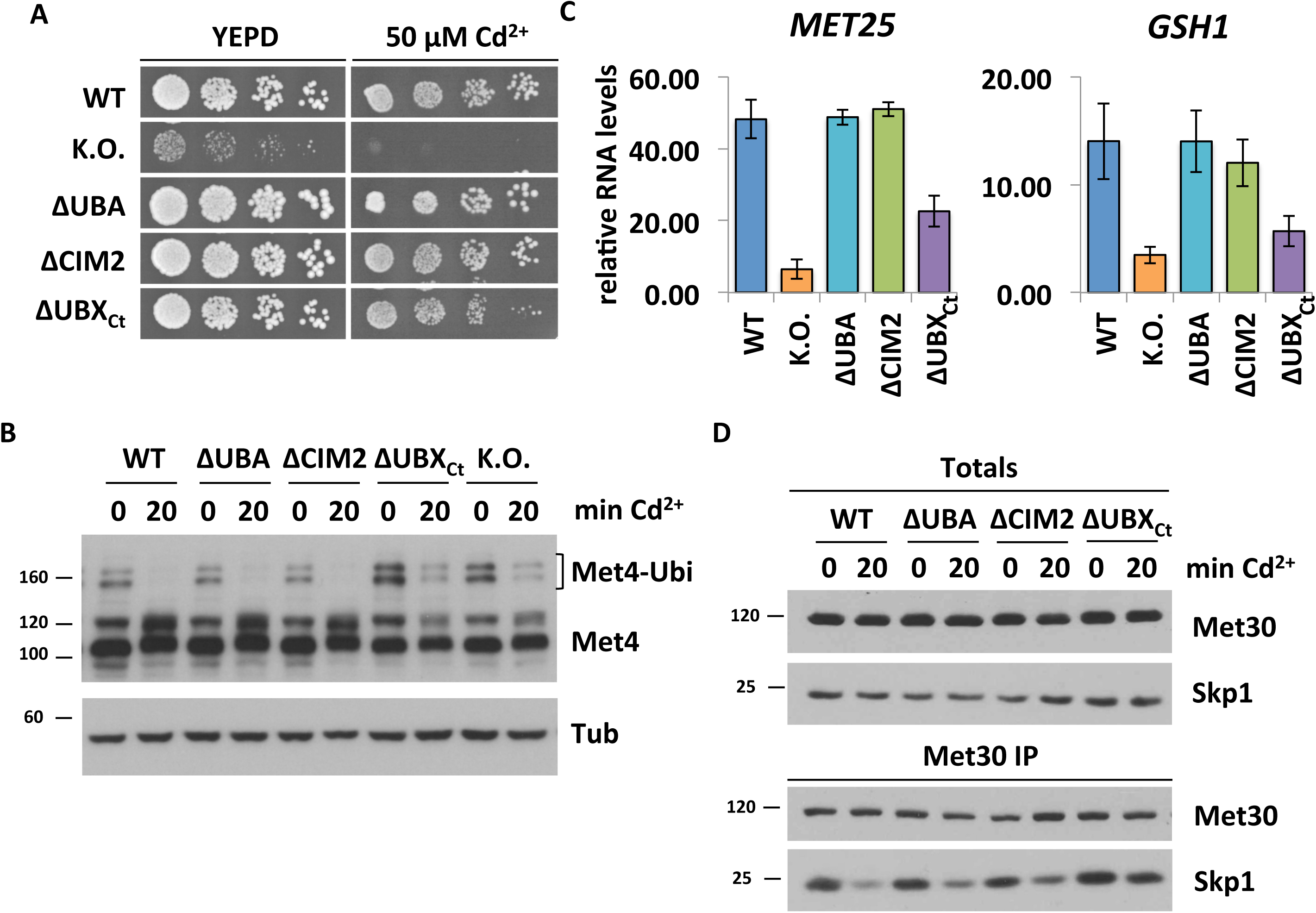
The UBX domain of Shp1 is required for SCF^Met30^ Disassembly during Cadmium Stress. A) *shp1-ΔUBX_Ct_* mutants are cadmium sensitive. Indicated strains were cultured to logarithmic growth phase, cells were counted and serial dilutions spotted onto YEPD plates supplemented with or without 50 µM CdCl_2_. Plates were incubated for two days at 30°C. B) The UBX domain is important for Met4 activation during heavy metal stress. Strains shown in Figure 3A were cultured at 30°C in YEPD medium and treated with 100 µM CdCl_2_ for 20 min. Whole cell lysates were analyzed by Western blotting using a Met4 antibody to follow the ubiquitylation and phosphorylation status of Met4. Tubulin was used as a loading control. C) Insufficient Met4 activation in *shp1-ΔUBX_Ct_* mutants is reflected in *MET25* and *GSH1* RNA levels. Strains as indicated were grown at 30°C in YEPD medium treated with 100 µM CdCl_2_ and samples were harvested after 40 min exposure. RNA was extracted and expression of Met4 target genes *MET25* and *GSH1* was analyzed by RT-qPCR and normalized to 18S rRNA levels (n=3). Data are represented as mean ±SD. D) *shp1ΔUBX_Ct_* mutants show altered dissociation kinetics during cadmium stress. ^12xMyc^Met30 of lysates shown in Figure 3B was immunoprecipitated and co-purified proteins were analyzed by Western blotting. Results shown in B and D are representative blots from three independent experiments.

These experiments cannot distinguish between a general requirement of Shp1 for Met4 activation and the cadmium signal specific pathway we propose. In addition to heavy metal stress methionine starvation blocks Met4 ubiquitylation and induces a similar transcriptional response program (25). However, Met4 activation by methionine stress is achieved through disruption of the Met30/Met4 interaction and not by SCF^Met30^ disassembly (16, 64). Importantly, Met4 activation in response to methionine starvation was unaffected in *shp1* mutants, demonstrating a specific requirement for Shp1 in the SCF^Met30^ disassembly pathway (Suppl. Fig 3A).

We next directly evaluated cadmium-induced SCF^Met30^ disassembly in *shp1* mutants. *shp1-Δubx_Ct_* mutants showed very slow Met30 dissociation, and the majority of Met30 remained in complex with the core ligase during heavy metal stress (Fig 3D). In contrast, all other mutants were indistinguishable from wild-type cells (Fig. 3D). In summary our results indicate, that an intact UBX domain of Shp1 is required for SCF^Met30^ disassembly during cadmium stress. Importantly, even though both ΔCIM2 and ΔUBX_Ct_ mutants showed severely decreased Shp1 interaction with Cdc48 (Fig 2C), only *shp1-Δubx_Ct_* mutants were deficient in SCF^Met30^ disassembly. Hence, the role of Shp1 in SCF^Met30^ during heavy metal stress is independent of a stable Cdc48-Shp1 interaction, but requires a specific contact between Cdc48 and Shp1 that is mediated by a contact interface at the very C-terminus of Shp1.

### Met30 Dissociation Kinetics during Heavy Metal Stress are Dependent on Shp1 Abundance

We next monitored Cdc48 and cofactor recruitment, as well as kinetics of SCF^Met30^ disassembly in the *shp1-Δubx_Ct_* mutant. Immunoprecipitated SCF^Met30^ formed a complex with Cdc48 and Shp1ΔUBX_Ct_ even under unstressed growth conditions (Fig 4A), further supporting that stable Cdc48-Shp1 binding is not required for recruitment of either. However, disassembly kinetics of SCF^Met30^ in *shp1Δ* and *shp1-Δubx_Ct_* mutants were severely delayed, demonstrating that physical separation of Met30 from the core SCF relies on Shp1, and specifically on its very C-terminal residues (Fig 4A, Suppl. Fig 4A). The slow disassembly of SCF^Met30^ in *shp1-Δubx_Ct_* mutants is also reflected in delayed Met4 activation indicated by persisting Met4 ubiquitylation (Fig. 4B). Surprisingly, similar to *shp1Δ* deletion mutants, the amount of Cdc48 in complex with Met30 under unstressed growth conditions was significantly increased in *shp1-Δubx_Ct_* mutants (Fig 4A; Suppl. Fig 4D). This increased steady state binding of Cdc48 to SCF^Met30^ in *shp1* mutants may be a result of the blocked dissociation and likely reflects accumulation of a dissociation intermediate (Fig 4A, Suppl. Fig 4C). Furthermore, even though Shp1-ΔUBX_Ct_ does not efficiently interact with Cdc48 (Fig. 2D), both Cdc48 and Shp1ΔUBX_Ct_ are efficiently recruited to SCF^Met30^ (Fig. 4A), indicating that both proteins are independently recruited to SCF^Met30^. However, due to the higher steady state binding of both Cdc48 and Shp1ΔUBX_Ct_ in *shp1-Δubx_Ct_* mutants, the fold-recruitment induced by cadmium stress was reduced for both proteins (Fig 4A, Suppl. Fig 4B). This decreased enrichment was not due to partial loss of Cdc48 binding, because Shp1ΔCIM2, which shows a similar reduction in Cdc48 binding (Fig. 2D), was recruited to SCF^Met30^ at similar levels as wild-type Shp1 (Suppl. Fig 4E). Importantly, despite the decreased fold-enrichment, the total amount of bound Cdc48 during cadmium stress is significantly higher in *shp1Δ* and *shp1-Δubx_Ct_* mutants than wild-type cells (Suppl. Fig 4D), likely a reflection of accumulation of the usually transient enzyme/substrate (SCF^Met30^/Cdc48) complex due to the blocked dissociation process. These results suggest that both Cdc48 and Shp1 are independently recruited to SCF^Met30^. However, their interaction through the C-terminal portion of the UBX domain on Sph1 is necessary to mediate Met30 dissociation.

**Figure 4.**
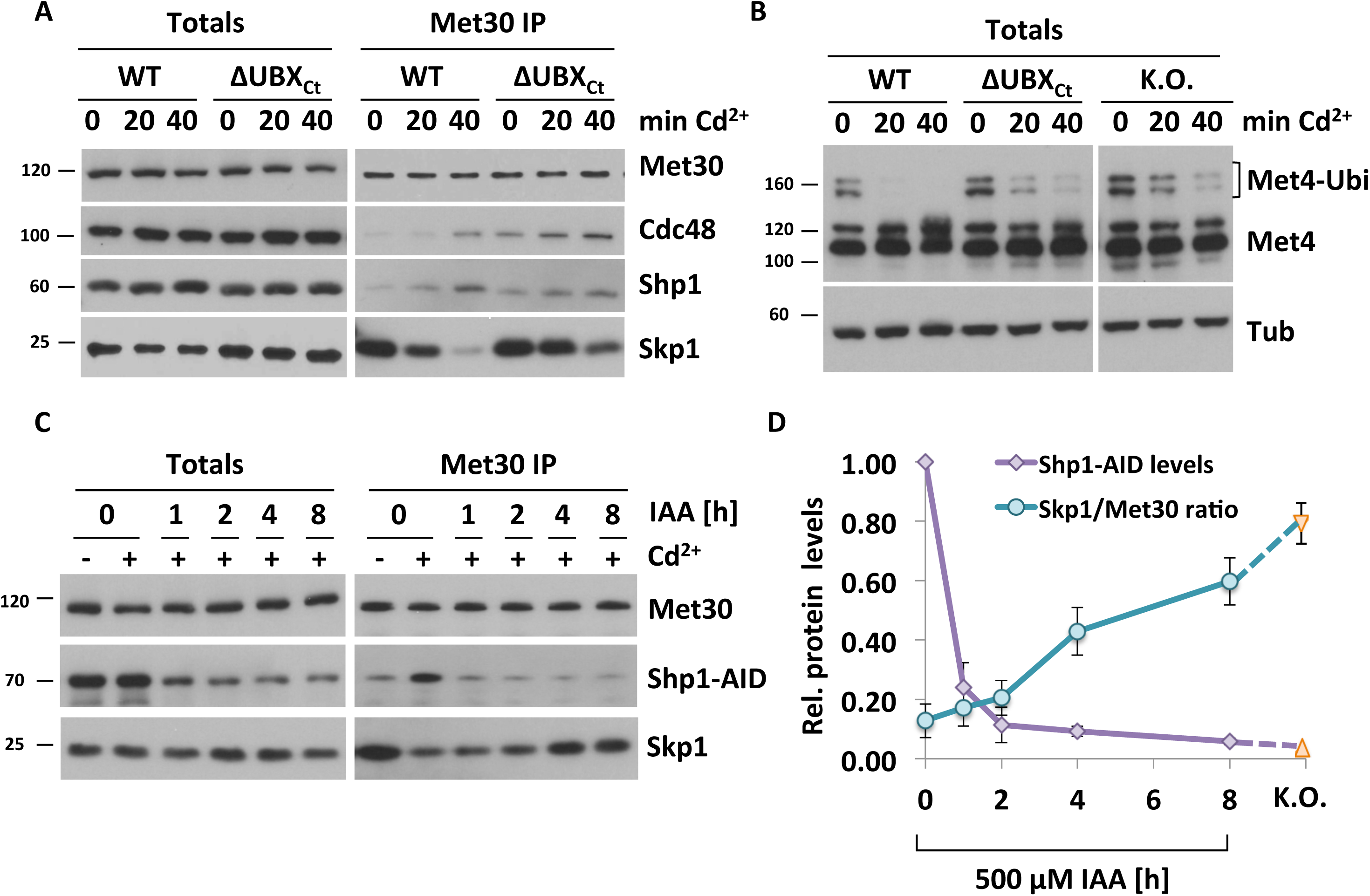
Met30 Dissociation during Cadmium Stress Depends on Shp1 Abundance. A) Cadmium-induced recruitment of Shp1 to SCF^Met30^ is significantly decreased in ΔUBX_Ct_ mutants. Strains expressing endogenous ^12xMyc^Met30, Cdc48^RGS6H^, Skp1, and Shp1^3xHA^, or ΔUBX_Ct_ Shp1^3xHA^ were cultured at 30°C in YEPD medium treated with 100 µM CdCl_2_ and samples were harvested at indicated time points. Whole cells lysates (Totals) were prepared and ^12xMyc^Met30 was immunoprecipitated and co-purified proteins were analyzed by Western blotting. B) *shp1-ΔUBX_Ct_* mutants show delayed Met4 activation. Whole cell lysates shown in Fig 4A were analyzed by Western blotting using a Met4 antibody to follow the ubiquitylation and phosphorylation status of Met4. Tubulin was used as a loading control. C) Met30 dissociation kinetics dependent on Shp1 abundance. Strains expressing endogenous ^12xMyc^Met30, Shp1^3xHA-AID^, and the F-Box protein ^2xFLAG^OsTir under the constitutive ADH promoter were cultured at 30°C in YEPD medium in the absence and presence of 500µM auxin for the indicated time to gradually down-regulate endogenous Shp1-AID levels. Cells were exposed to 100 µM CdCl_2_ and samples were harvested after 20 min. ^12xMyc^Met30 was immunoprecipitated and co-purified Skp1 and Shp1-AID were analyzed by Western blotting. Results shown in A, B and C are representative blots from three independent experiments. D) Densitometric analysis of Figure 4C shows a correlation between Shp1-AID abundance and Met30 dissociation from the SCF core during cadmium stress. Purple line = densitometric analysis of Western blot band intensities of Shp1-AID Totals. Time point 0 was set to 1. Turquoise line = co-immunoprecipitated Skp1 signals were normalized to immunoprecipitated ^12xMyc^Met30 signals to quantify F-box protein dissociation from the core ligase after 20 min of cadmium exposure. Orange triangles correspond to the values obtained from Fig 1A and Suppl. Fig 1A quantifications of *shp1Δ* mutants.

The human Shp1 homolog p47 has a concentration dependent effect on Cdc48 activity (57). We therefore investigated if SCF^Met30^ bound Shp1 levels correlated with Met30 dissociation during cadmium stress, and whether blocked disassembly in *shp1-Δubx_Ct_* mutants was due to reduced Shp1ΔUBX_Ct_ in the dissociation complex. The AID system enabled us to titrate Shp1 levels, ranging from endogenous to about ten-fold lower levels (Fig 4C). Met30 dissociation was attenuated when Shp1-AID levels were decreased, revealing a correlation between Shp1 abundance and cadmium-induced SCF^Met30^ disassembly (Fig 4D, Suppl. Fig 4F). A significant dissociation defect was only observed, when Shp1-AID levels were reduced by more than 75%. Importantly, Shp1ΔUBX_Ct_ amounts recruited to the SCF^Me30^/Cdc48 complex are significantly above the Shp1 threshold level necessary for complete SCF^Me30^ disassembly (Fig 4A, Fig. 4C, 1h IAA, Suppl. Fig 4H). Thus, the defect of Met30 dissociation observed in *shp1-Δubx_Ct_* mutants is not due to reduced Shp1ΔUBX_Ct_ recruitment, but reflects requirement of UBX mediated interaction with Cdc48 to stabilize the complex and trigger the dissociation process.

## Discussion

Cdc48 is an exceptionally versatile protein and substrate specificity is thought to depend on cofactors (65). During heavy metal stress Cdc48 plays a critical role in signal-induced disassembly of the SCF^Met30^ ubiquitin ligase complex (16). Our results show that the UBA-UBX cofactor Shp1 is critical for this process. Like Cdc48, Shp1 is recruited into SCF^Met30^ upon cadmium exposure (Fig1A, Suppl. Fig 1A). Interestingly, the prominent heterodimeric Ufd1/Npl4 cofactor pair also gets recruited to Met30 under heavy metal stress conditions (Suppl. Fig 5 A&B). However, temperature-sensitive *ufd1-2* and *npl4-1* mutants do not exhibit cadmium sensitivity, Met4 activation defects, or a dissociation phenotype during heavy metal stress (16). Likely cadmium-induced recruitment of Ufd1/Npl4 reflects their canonical function in this context and is necessary for targeting dissociated Met30 for degradation by the proteasome. Cadmium stress generates an excess of ubiquitylated “Skp1-free” Met30, which is rapidly degraded by a non-canonical pathway via a Cdc53-dependent, but Skp1-independent proteolytic pathway (66). This pathway is necessary to prevent substrate shielding by Met30 that is separated from SCF (66). Consistent with this model, Met30 that cannot associate with Skp1 due to deletion of the F-box motif (Met30^ΔF-box^), is stabilized in temperature-sensitive *ufd1-2* and *npl4-2* mutants (Suppl. Fig. 5 C). Therefore, we speculate that there might be two populations of Cdc48 complexes involved in processing of SCF^Met30^ during cadmium stress. The Cdc48^Shp1^ complex is required for the disassembly of the SCF^Met30^complex, and Cdc48^Ufd1/Npl4^ for degradation of unbound Met30 via the proteasomal pathway.

Surprisingly, Met30 protein levels were significantly decreased in *shp1*Δ mutants compared to wild-type cells (Fig 1A, Fig 2B). While Shp1 is involved in the degradation of a Cdc48-dependet model substrate (67), the half-life of Met30 is unaffected in *shp1Δ* cells and all other *shp1* mutant variants we tested (Suppl. Fig 2F). However, *MET30* RNA levels were significantly decreased in the *shp1Δ* knock out strain (Suppl. Fig 2G). Accordingly, low Met30 protein levels are due to a transcriptional defect. Since all other Shp1 mutants showed normal RNA levels, we hypothesize that altered transcription might be associated with other aspects of Shp1 function (55).

Closer examination of functional domains in Shp1 revealed several interesting findings. We expected that the UBA domain with its ubiquitin binding ability plays a role in Shp1 recruitment to Met30, which is autoubiquitylated during heavy metal stress (68). Additionally, it has been proposed that UBA domains can inhibit ubiquitin chain elongation (69). However, deletion of the UBA domain in Shp1 did not show any cadmium specific phenotypes and is not required for SCF^Met30^ or Met4 regulation (Fig 3).

It was also unexpected that Cdc48 and Shp1 were independently recruited to SCF^Met30^ upon cadmium exposure (Fig 1A & Fig 3A, Suppl. Fig 4E), because Shp1 is believed to be constitutively bound to Cdc48 to fulfill its cellular tasks (46, 51, 53). While independent recruitment of Cdc48 is supported by results in cells lacking Shp1 (Fig. 1A), autonomous recruitment of Shp1 was suggested based on experiments using mutant versions of Shp1 with dramatically reduced Cdc48 steady-state binding activity. We can therefore not completely exclude that under normal conditions Shp1 is co-recruited as a Cdc48-associated cofactor. However, we can conclude that independent binding sites for Cdc48 and Shp1 exist on SCF^Met30^ that drive recruitment. Furthermore, the interaction of Shp1ΔCIM2 with Cdc48 is dramatically compromised (Fig 2D), yet Shp1ΔCIM2 is efficiently recruited to SCF^Met30^ and *shp1-Δcim2* mutants disassemble and effectively inactivate SCF^Met30^ indistinguishable from wild-type cells (Fig 3, Suppl. Fig 4E). These findings further support two independent binding sites for Cdc48 and Shp1 on SCF^Met30^, and the proposed model that mechanical transmission of forces to separate Met30 from the core SCF requires at least two anchor points.

Truncation of the last 22 residues of Shp1 (ΔUBX_Ct_), like mutation of the CIM2 domain, results in severely reduced steady state binding to Cdc48 (Fig 2D). However, in contrast to *shp1-Δcim2* mutants, *shp1-ΔUBX_Ct_* mutants fail to efficiently disassemble SCF^Met30^ and activate Met4 (Fig 3). The reduced interaction with Cdc48 cannot be the reason for failure to dissociate Met30, because Shp1ΔCIM2 is equally deficient in Cdc48 interaction, yet recruitment to SCF^Met30^ and Met30 dissociation is undistinguishable from wild-type cells (Suppl. Fig 4E). Our results more likely suggest that the C-terminus of the UBX domain is important for proper topological assembly of a functional disassembly complex to stimulate the Cdc48 ATPase activity and generate the required force for Met30 dissociation. In addition, the residues we deleted are potential posttranslational modification sites in the human orthologue p47 (70, 71). Lack of these modifications might hinder correct interactions of the disassembly complex.

This necessity for the 22 C-terminal residues of Shp1 to trigger disassembly but dispensability for recruitment presents as a separation of function mutant, that allows formation but not activation of the disassembly complex. Consequently, we observed accumulation of Cdc48 on SCF^Met30^ in *shp1-ΔUBX_Ct_* mutants, which likely presents a dissociation intermediate, trapped just before the step of SCF^Met30^ disassembly. Surprisingly, even during normal, unstressed growth conditions, the usually very low amount of Cdc48 in complex with Met30 is significantly increased in *shp1-ΔUBX_Ct_* mutants (Fig 4A), suggesting continuous low-level dissociation of Met30 from the core SCF. This process may be part of a quality control mechanism that monitors proper assembly of the ubiquitin ligase complex.

While Cdc48/p97 can bind to ubiquitin directly *in vitro* (31). UBX cofactors are thought to be adaptors that regulate the interaction between Cdc48 and ubiquitylated substrates (46). Under normal growth conditions Met30 is ubiquitylated at low levels, but cadmium stress drastically increases Met30 autoubiquitylation (68). This autoubiquitylation is the recruitment signal for Cdc48 and given that Cdc48 recruitment is independent of Shp1 and Npl4/Ufd1 (Fig 1A, Fig3A) (16), a direct interaction with ubiquitylated Met30 is likely. However, Shp1 is required to properly orient the complex, activate the ATPase activity, and provide a second attachment point for transmission of the mechanical force.

Initially, we used the AID system for induced down regulation of Shp1 to study acute loss of Shp1 function and eliminate possible indirect effects caused by secondary mutations acquired by the *shp1Δ* deletion strain. However, it also became a handy tool to closer examine dependence of SCF^Met30^ disassembly on different levels of Shp1. This question was interesting because different concentration-dependent binding modes of p47 to p97 have been suggested to control p97 ATPase activity (57). Decreased Shp1-AID levels resulted in impaired SCF^Met30^ dissociation (Fig 1D), and titration of Shp1-AID levels uncovered a correlation between Shp1 abundance and disassembly (Fig 4 C&D). This phenomenon is likely related to a need for a minimum concentration of Shp1 in the dissociation complex to activate Cdc48. When first discovered, human p47 was described as an inhibitor of p97 activity (72) However, a later study showed a more complex mechanism of ATPase regulation by p47, in which higher concentrations of p47 bound to p97 relieve inhibition of p97 ATPase activity (57). The authors suggest, that a transition from monomeric to trimeric p47 on p97 is responsible for this effect (57). We hypothesize that at low amounts, Shp1 facilitates maximum inhibition of Cdc48 ATPase activity. Upon cadmium exposure, Shp1 is recruited to SCF^Met30^, increasing the local concentration of the cofactor. This increase might lead to Shp1 trimerization and stimulation of ATP hydrolysis followed by separation of Met30 from the core SCF. Future experiments will need to address this hypothesis.

In summary, our study identifies the UBA-UBX family member Shp1 as the key cofactor for signal induces disassembly of SCF^Met30^. We show that the ATPase Cdc48 and Shp1 are recruited independently to SCF^Met30^ during cadmium stress, but interaction through the C-terminal end of the Shp1 UBX domain is necessary to activate the dissociation complex and catalyze separation of Met30. These results provide insight into ubiquitin-dependent, signal-induced, active remodeling of multi-protein complexes to control their activity.

## Acknowledgments

We thank S. Jentsch, T. Rapoport, P. Silver, and R. Hampton for yeast strains; R. Deshaies, W. Harper, M. Tyers, T. Sommer, and E. Jarosch for antibodies.

This work was supported by the National Institute of Health grant R01 GM-066164 to P.K. and the Hitachi-Nomura Award to L. L..

## Supplemental Information

**Supplementary Figure 1.**
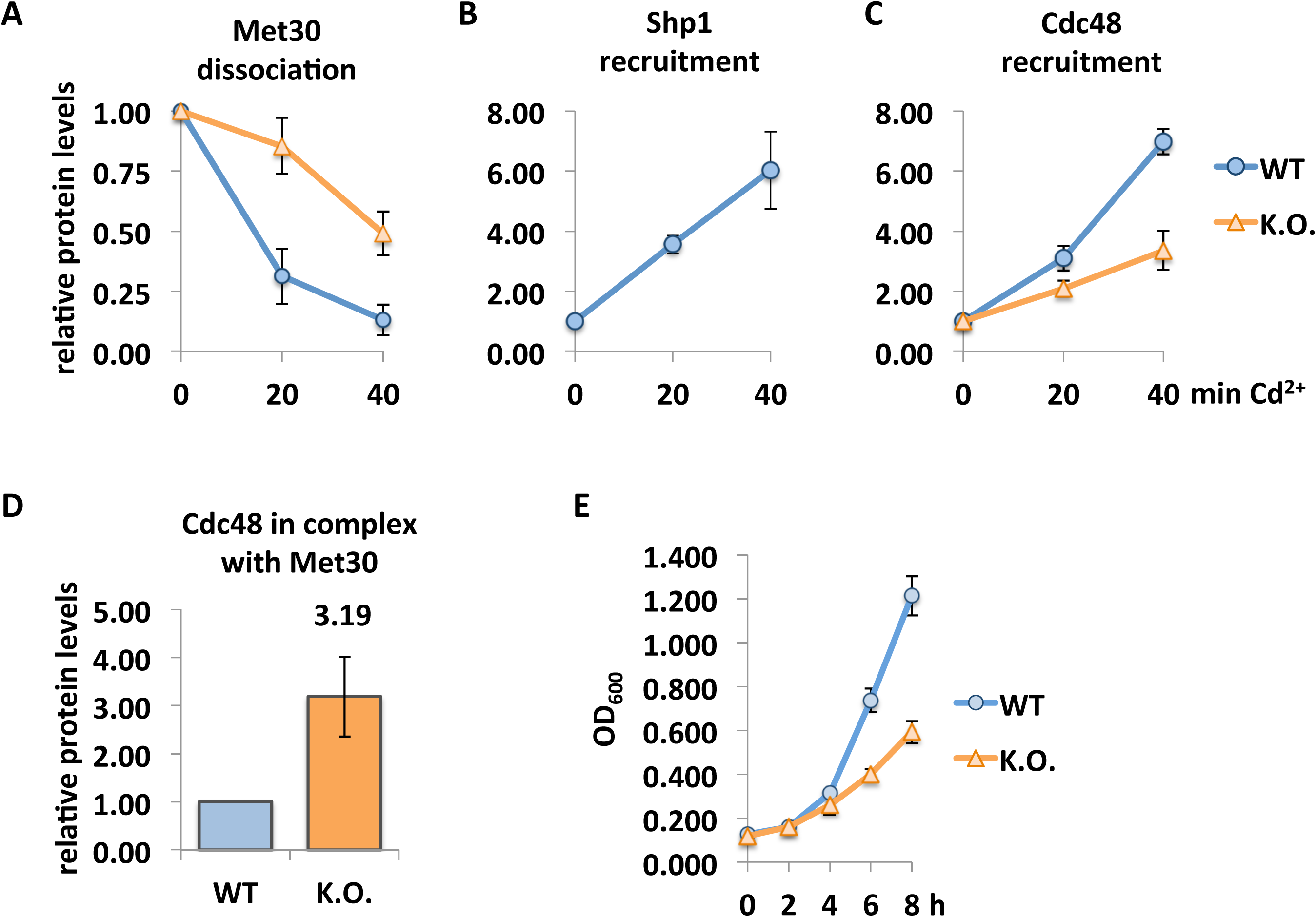
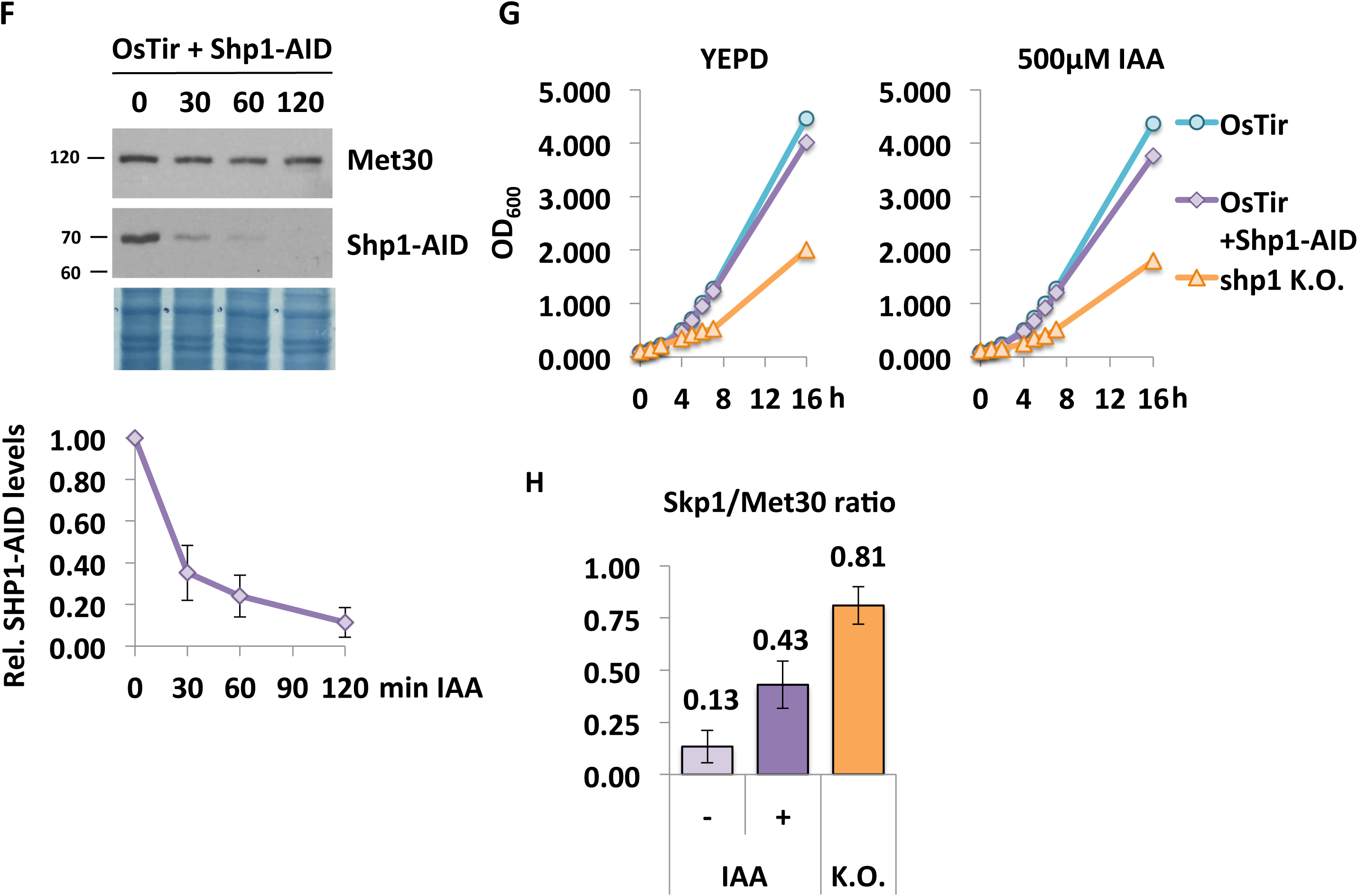
Growth curves, dissociation and recruitment kinetics in WT, Shp1 knock out and Shp1-AID strains. A-C) Densitometric analysis of Western blot band intensities of immunoprecipitations in Fig 1A. For quantifications in A-C the signal intensities for each Shp1 variant at time point 0 were set to 1. A) Skp1 signals were normalized to ^12xMyc^Met30 to quantify Met30 dissociation from the core ligase. B) Shp1^3xHA^ signals were normalized to ^12xMyc^Met30 to determine recruitment of Shp1 to SCF^Met30^. Cdc48^RGS6H^ signals were normalized to ^12xMyc^Met30 to resolve recruitment into SCF^Met30^. D) Densitometry of Cdc48 ^RGS6H^ /^12xMyc^Met30 ratios, WT at time point 0 was set to 1. E) *shp1Δ* mutants show a significant growth defect at 30°C. WT, and Shp1 K.O. strains were grown at 30°C in YEPD medium and samples were taken at indicated time points. Optical density was measured at 600nm. F) Shp1-AID gets sufficiently down-regulated upon auxin exposure. Strains expressing endogenous ^12xMyc^Met30, Shp1^3xHA-AID^ and the F-Box protein ^2xFLAG^OsTir under the constitutive ADH promoter were cultured at 30°C in YEPD. Samples were taken at indicated time-points after the addition of 500µM auxin (IAA). Protein levels were analyzed by Western blotting. The amido black stain of the membrane is shown as a loading control. G) Depicted strains do not show a significant growth defect at 30°C in the absence or presence of auxin. Strains were grown at 30°C in YEPD medium or YEPD containing 500 µM IAA and samples were taken at indicated time points. Optical density was measured at 600nm. H) Densitometric analysis of Western blot band intensities of immunoprecipitations in Fig 1D. Skp1 signals were normalized to ^12xMyc^Met30 to determine Met30 dissociation from the core ligase during cadmium stress.

**Supplementary Figure 2.**
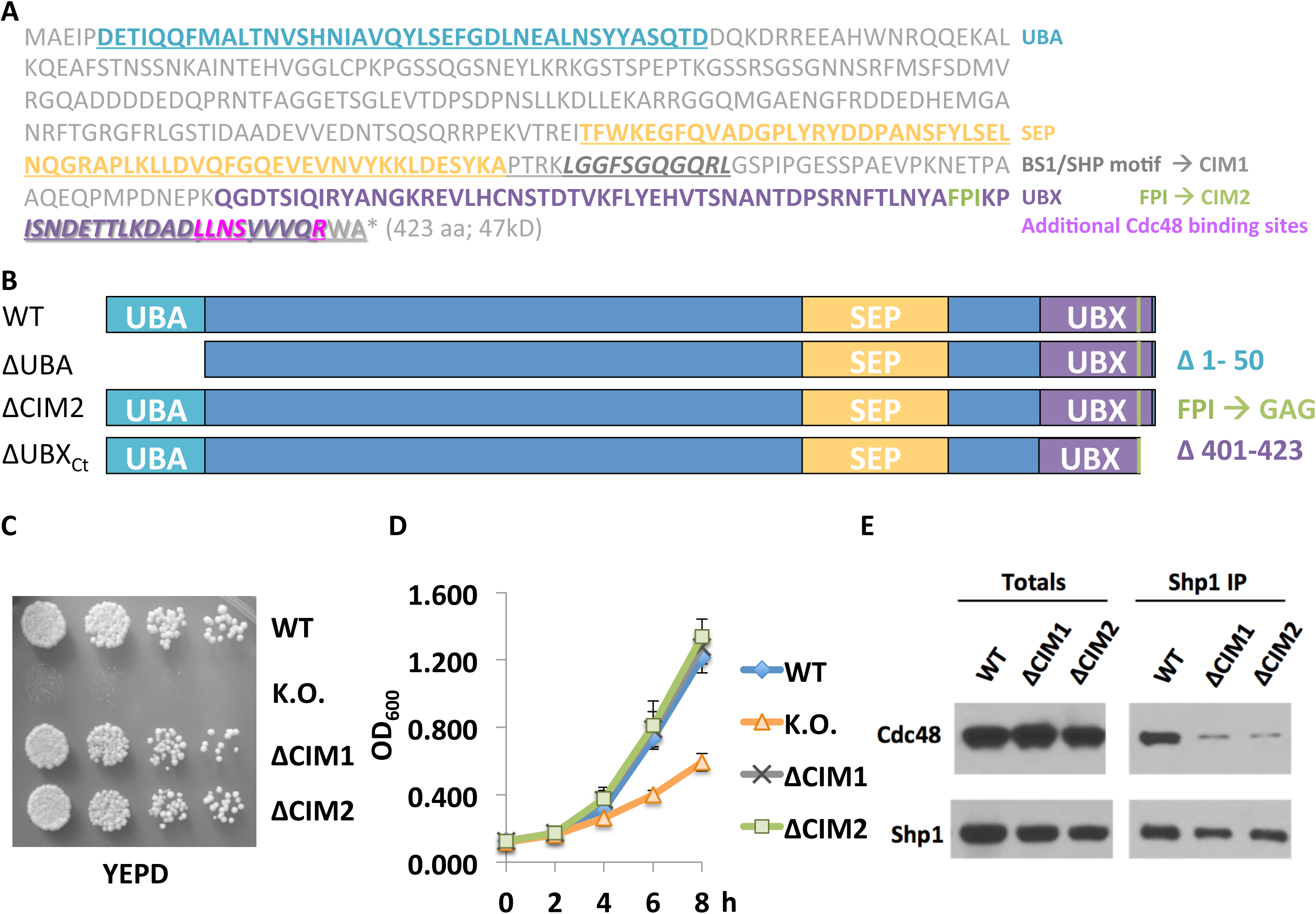
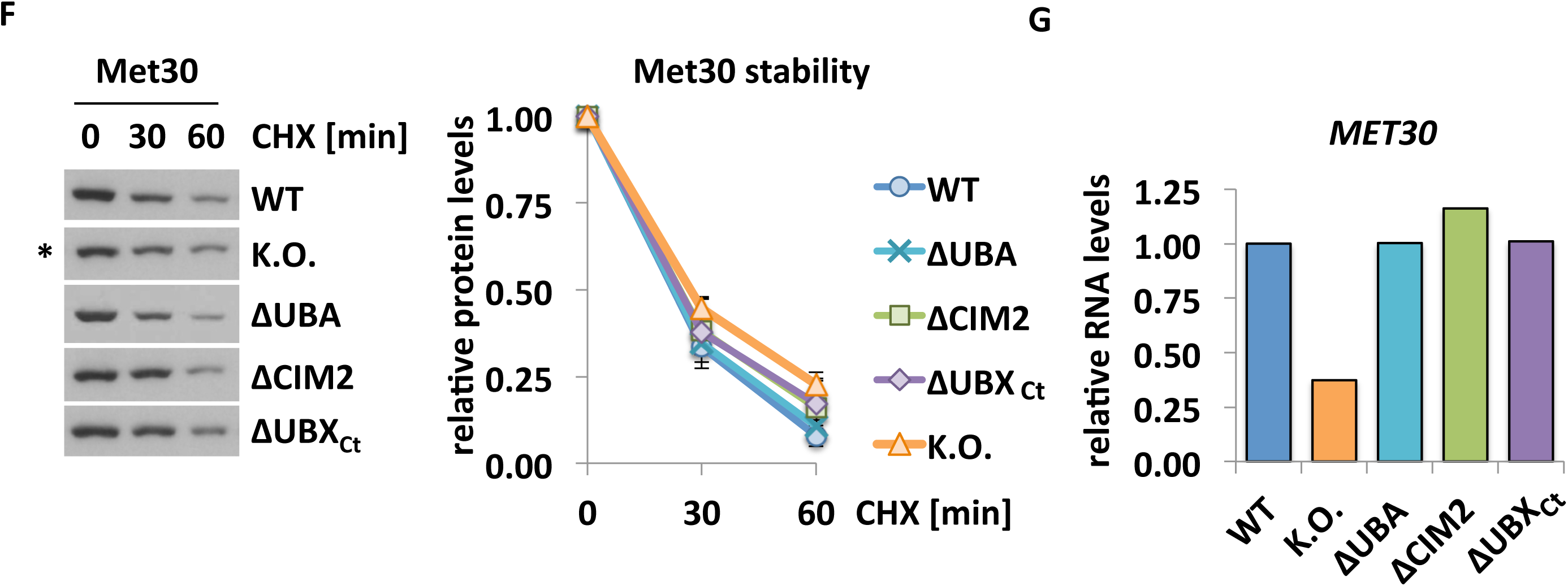
A) Amino acid sequence of Shp1 - Specific domains/motifs are labeled and color-coded. B) Schematic of the most important Shp1 variants used throughout this study. C) Spotting assay - Serial dilutions of depicted strains were made and spotted onto YEPD plates. Plates were incubated for two days at 30°C. D) Shp1 WT, ΔCIM1 &2 and K.O. strains were grown at 30°C in YEPD medium and samples were taken at indicated time points. Optical density was measured at 600nm. E) Cdc48 binding is significantly decreased in ΔCIM1 and ΔCIM2. SHP1^3xHA^ variants were immunoprecipitated and co-purifications of Cdc48^RGS6H^ were analyzed by Western blotting. F) Met30 protein stability is unaffected in *shp1Δ* deletion mutants. Depicted strains were grown at 30°C in YEPD medium, cycloheximide (100 µg/ml) was added, and samples were collected at indicated time points. Protein stability was analyzed by immunoblotting followed by densitometric analysis. Asterisk next to Met30 levels of K.O. (*shp1Δ*) indicates longer exposure of the Western blot panel to show similar starting amounts of Met30 in CHX time course. G) *MET30* RNA levels are decreased in *shp1Δ* cells. Depicted cultured were grown at 30°C in YEPD medium. RNA was extracted and expression of *MET30* was analyzed by RT-qPCR and normalized to 18S rRNA levels.

**Supplementary Figure 3.**
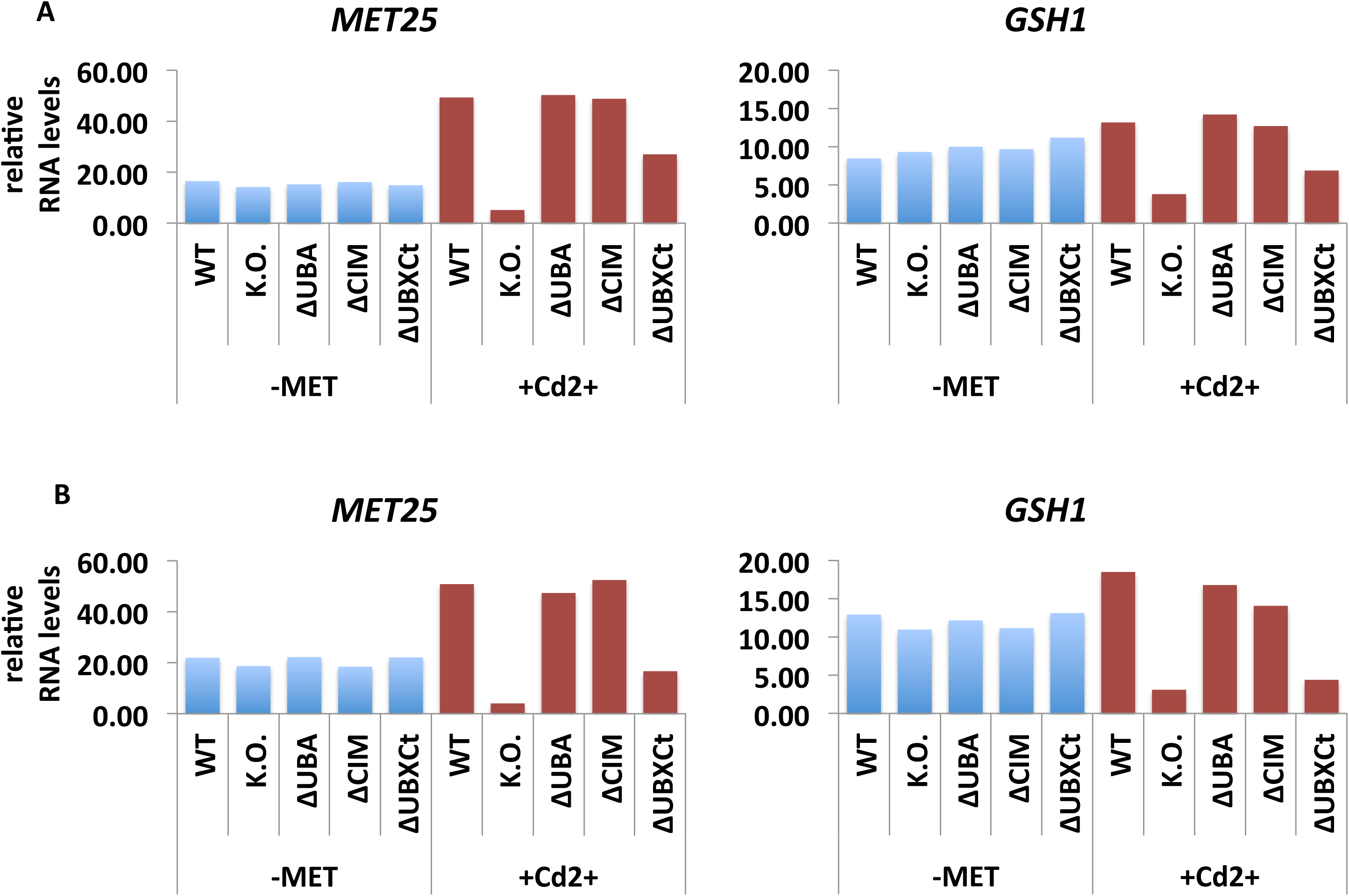
A&B) The expression of Met4-dependent genes in response to methionine starvation. Depicted yeast strains were grown at 30°C in YEDP medium to OD_600_ of 0.6. For Methionine starvation, cultures were washed with water and shifted to minimal medium without methionine for 30 minutes and then harvested. For heavy metal stress induction, cultures were treated with 100µM CdCl_2_ and samples were harvested after 40 min exposure. RNA was extracted and expression of Met4 target genes *MET25* and *GSH1* was analyzed by RT-qPCR and normalized to 18S rRNA levels (n=2). Set I shown in A, set II shown in B.

**Supplementary Figure 4.**
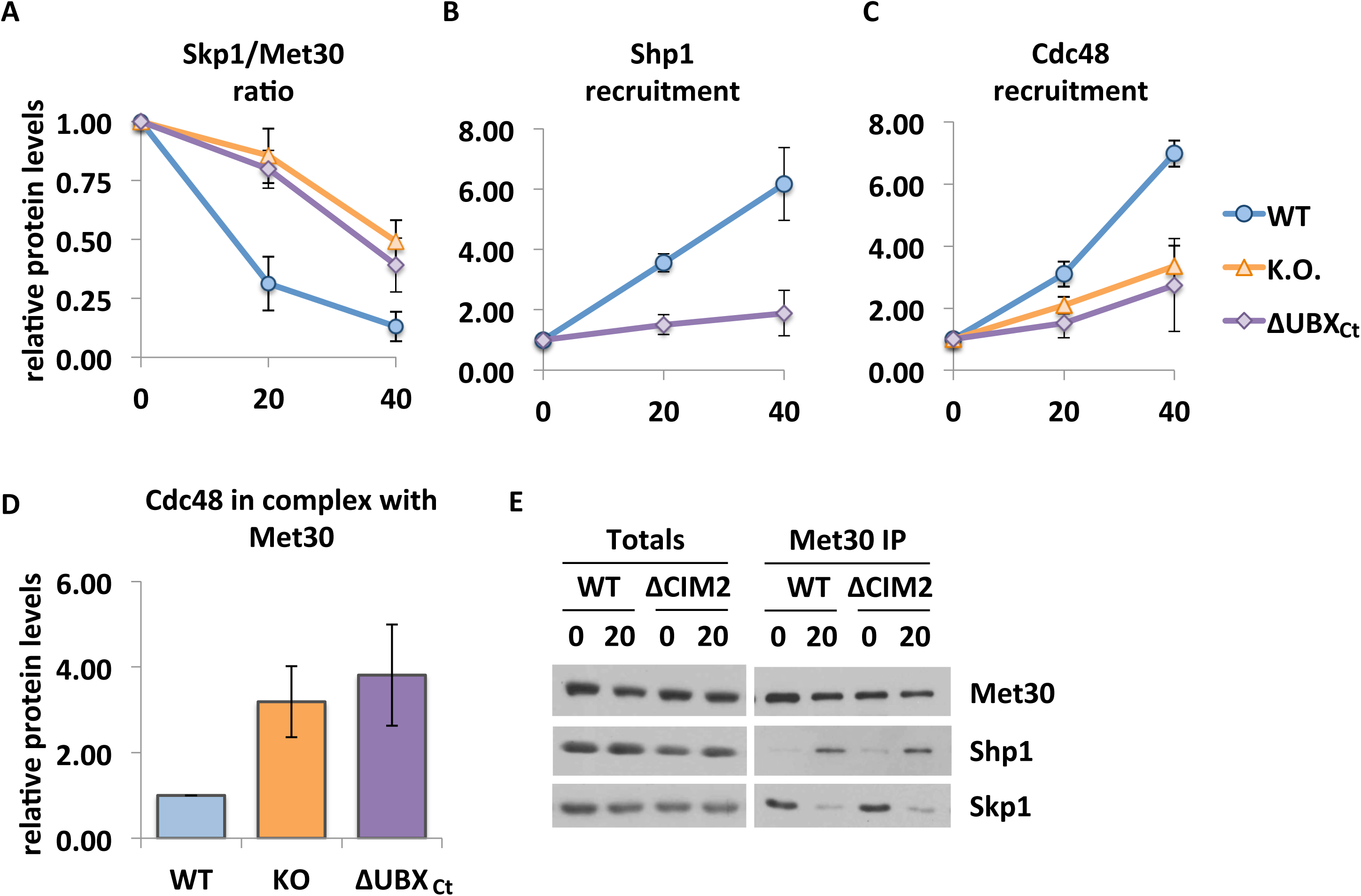
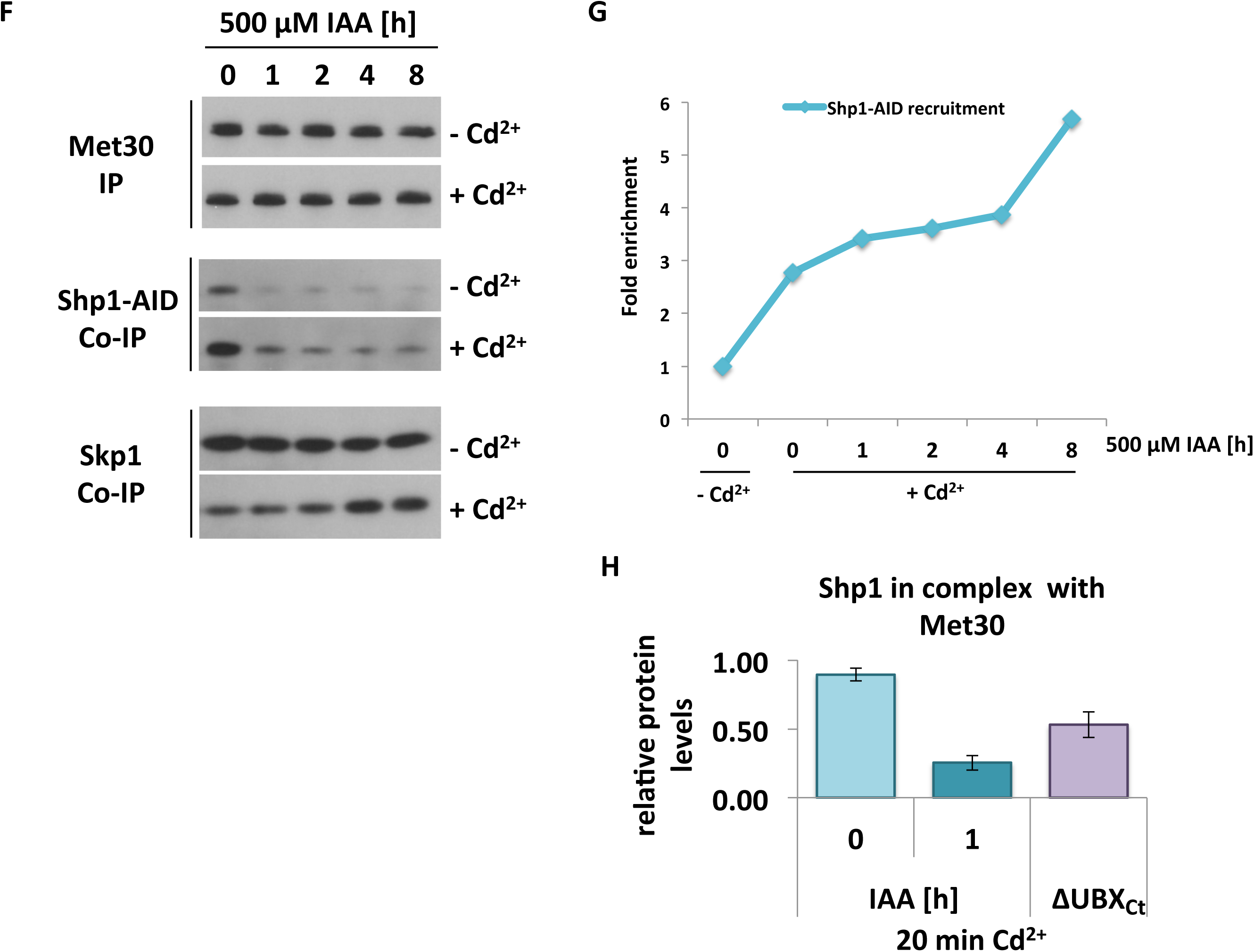
A-D) Quantification data of ΔUBX_Ct_ was integrated in the graphs shown in Suppl. Fig 1 A-D. For quantifications in A-C the signal intensity of each Shp1 variant at time point 0 was set to 1. A) co-immunoprecipitated Skp1 signals were normalized to immunoprecipitated ^12xMyc^Met30 to quantify Met30 dissociation from the core ligase. B) Shp1^3xHA^ signals were normalized to ^12xMyc^Met30. Then the ratio of co-immunoprecipitated Shp1^+Cd^/Shp^w/o Cd^ was determined to analyze recruitment of Shp1 to SCF^Met30^. Cdc48^RGS6H^ signals were normalized to ^12xMyc^Met30 and the ratio of co-immunoprecipitated Cdc48^+Cd^/Cdc48^w/o Cd^ was analyzed to resolve recruitment of Cdc48 to SCF^Met30^. D) Densitometry of co-immunoprecipitated Cdc48 ^RGS6H^/ immunoprecipitated ^12xMyc^Met30 ratio. WT at time point 0 was set to 1. E) Shp1ΔCIM2 is recruited to SCF^Met30^ during cadmium stress. WT and *shp1ΔCIM2* mutants were cultured at 30°C in YEPD medium and treated with 100 µM CdCl_2_ and samples were harvested after 20 min of exposure. ^12xMyc^Met30 was immunoprecipitated and co-precipitated proteins were analyzed by Western blotting. F) Full panel of immunoprecipitations that was partially shown in Fig 4C. A strain expressing endogenous ^12xMyc^Met30, Shp1^3xHA-AID,^ and the F-Box protein ^2xFLAG^OsTir under the constitutive ADH promoter was cultured at 30°C in YEPD medium in the absence and presence of auxin for the indicated time to gradually down-regulate endogenous Shp1-AID levels. Cells were exposed to 100µM CdCl_2_ and samples were harvested after 20 min. ^12xMyc^Met30 was immunoprecipitated and co-purified Skp1 was analyzed by Western blotting. G) Densitometric analysis of Western blot band intensities of immunoprecipitations in Suppl. Fig 4F. Shp1^3xHA^ signals were normalized to ^12xMyc^Met30. Then the ratio of co-immunoprecipitated Shp1^+Cd^/Shp^w/o Cd^ was determined to follow Shp1 recruitment. H) Densitometry of Shp1^3xHA^/^12xMyc^Met30 ratio. Quantifications of Western blots shown in Fig. 4B and 4C.

**Supplementary Figure 5.**
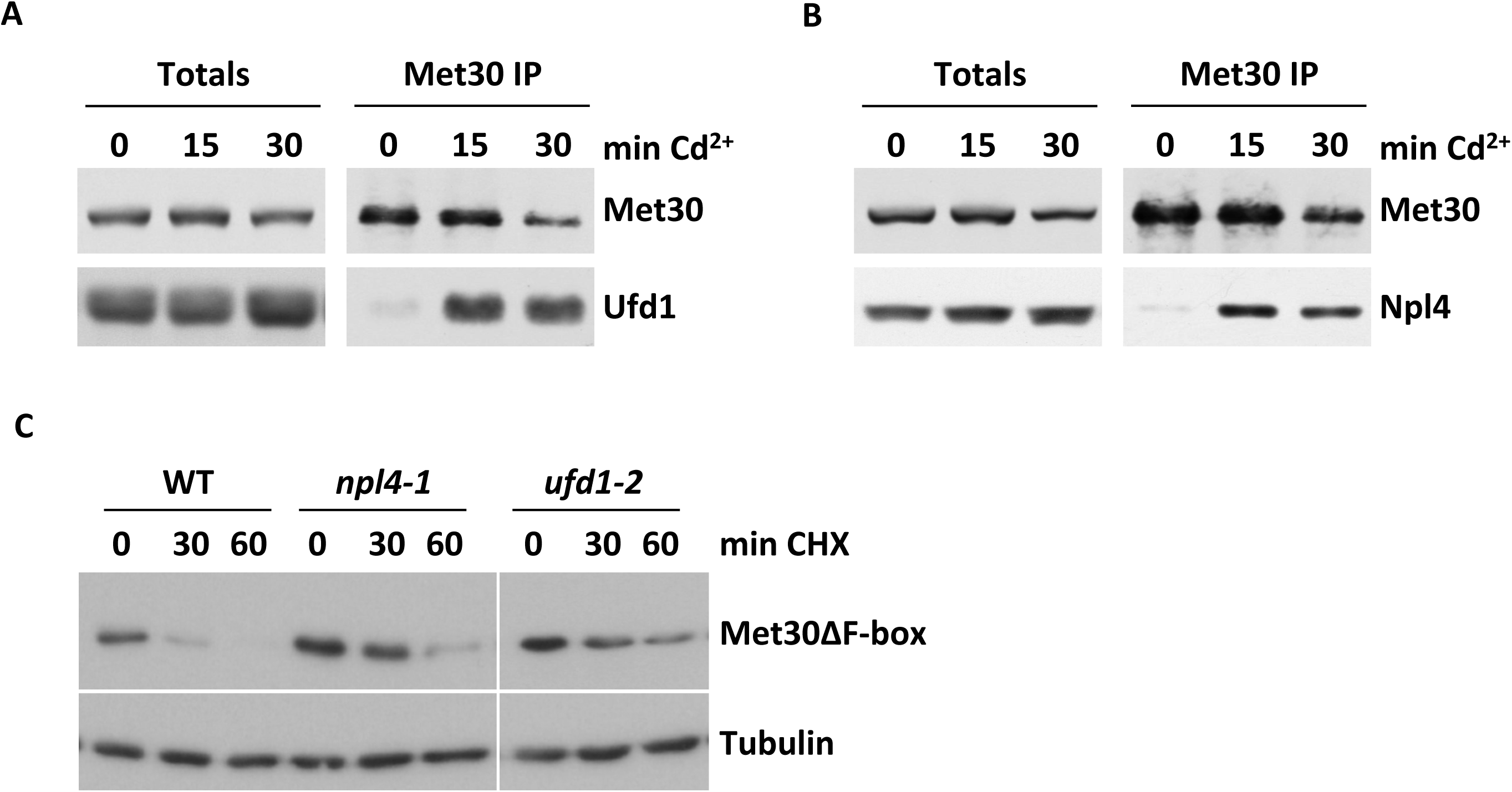
A&B) Cdc48 co-factors Ufd1 and Npl4 are recruited to SCF^Met30^ during heavy metal stress. Strains expressing endogenous ^12xMyc^Met30, Skp1, and Ufd1^3xHA^ or Npl4^3xHA^ respectively were cultured at 30°C in YEPD medium, treated with 100 µM CdCl_2_, and samples were harvested at indicated time points. ^12xMyc^Met30 was immunoprecipitated and co-precipitated proteins were analyzed by Western blotting. C) Cdc48 cofactors Npl4 and Ufd1 are involved in ‘Skp1-free’ Met30 degradation. Wild type, *npl4-1*, and *ufd1-2* temperature sensitive strains expressing ^12myc^Met30ΔFbox were cultured at permissive temperature (25°C) and then shifted to non-permissive temperature (37°C) for 1.5 h. Cycloheximide (100 µg/ml) was added and samples were collected at the time intervals indicated. Met30ΔFbox protein stability was analyzed by immunoblotting with anti-myc antibodies. Tubulin was used as a loading control.

## Methods

### Plasmids, Yeast Strains and Growth Conditions

Yeast strains used in this study are isogenic to 15DaubD, a bar1D ura3Dns, a derivative of BF264-15D (1). Specific strains are listed in Table S1. The 9xMYC tag of pADH_OsTir9xMYC::URA (2) was exchanged with a 2xFLAG tag by amplification of the vector using primers 1&2 (table S3). The PCR fragment was digested with BamHI and ligated to generate pADH_OsTir2xFLAG::URA. The plasmid was linearized with StuI and transformed into yeast strain PY1. Next, The AID-fragment was amplified from the pKAN-pCUP1-9xMYC-AID plasmid using primers 3&4 (table S3) and inserted via Gibson assembly into the AscI linearized pFA6a-3xHA::TRP plasmid to generate the tagging vector pFA6a-3xHA-AID::TRP. A PCR fragment using primers 13&14 (table S3) was amplified and transformed into PY1+pADH_OsTir2xFLAG::URA. This strain was transformed with YCpLEUpmet30_9xMYCMET30 to generate PY2230. For CRISPR/Cas9 edited yeast strains the vector pML107 (3) which contain both sgRNA and Cas9 expression cassettes was used. To generate the guide RNA sequences the “CRISPR Toolset” (http://wyrickbioinfo2.smb.wsu.edu/crispr.html) was consulted. pML107 was linearized using SwaI and hybridized primers for gRNAs as shown in table S3 were inserted via Gibson assembly to generate pML107-Shp1 vectors (table S2). The vectors were transformed together with the hybridized 90mer oligos containing the Shp1-specific repair sequence into PY1630 to generate marker-free Shp1::3xHA mutants shown in table S1. All other strains used were generated using classic PCR based gene modification methods (4). Generated plasmids and strains were verified by sequencing. All strains were cultured in standard media, and standard yeast genetic techniques were used unless stated otherwise (5). References to the use of cadmium (Cd^2+^) are specifically to cadmium chloride (CdCl_2_). For cell spotting assays, strains were cultured to logarithmic growth phase, sonicated and then counted. Serial dilutions were made and spotted onto YEPD plates supplemented with or without indicated amounts of CdCl_2_via a pin replicator (V&P Scientific, San Diego, CA).

### Protein Analysis

For Western blot analyses and immunoprecipitation assays, yeast whole cell lysates were prepared under native conditions in Triton X-100 buffer (50mM HEPES, pH 7.5, 0.2% Triton X-100, 200mM NaCl, 10% glycerol, 1mM dithiothreitol, 10mM Na-pyrophosphate, 5mM EDTA, 5mM EGTA, 50mM NaF, 0.1mM orthovanadate, 1mM phenylmethylsulfonyl floride [PMSF], and 1mg/ml each leupeptin, and pepstatin). Cells were homogenized in a screw-cap tube with 0.5 mm glass beads and Antifoam Y-30 using a MP FastPrep 24 (speed 4.0, 3×20 sec). Lysates were separated from glass beads and transferred into 1.5 ml reaction tubes (USA Scientific). Lysates were cleared by centrifugation (10 min, 14000g at 4°). For Western blot analyses proteins were separated by SDS-PAGE and transferred to a polyvinylidene difluoride (PVDF) membrane. Proteins were detected with the following primary antibodies: anti-Met4 (1:20000; a gift from M. Tyers), anti-Skp1 (1:5000; a gift from R. Deshaies), anti-Cdc48 (1:20000; a gift from E. Jarosch), anti-Myc (1:2000; Santa Cruz, 9E10), anti-HA (1:2000; Santa Cruz, F7) anti-RGS6H (1:2000; QIAGEN, Germantown, MD) anti-Tubulin (1:2000; Santa Cruz). Western blots results shown are representative blots from three experiments with independent cultures. Band intensities of immunoblots were quantified using either ImageJ or the Biorad Image Lab software. For immunoprecipitation of 9xMYC tagged proteins, 1 mg of total protein lysate was incubated in Triton X-100 buffer equilibrated MYC-trap beads (Chromotek) in a final volume of 500 µl over night at 4°C on a nutator. The next day beads and supernatants were separated by centrifugation (2 min, 100g at 4°C). Beads were washed 3 times in 1 ml Triton X-100 buffer plus inhibitors at 4°C. Beads were resuspended in 2x Laemmli buffer and boiled for 5 min to elute proteins. For immunoprecipitation of 3xHA tagged proteins, 1 mg of total protein lysate in a final volume of 500 µl was incubated with 1 µg of anti-HA Y-11 (Santa Cruz, discontinued) for 2 hours at 4°C on a nutator. Triton X-100 buffer equilibrated protein A sepharose (Sigma-Aldrich) was added and incubated over night at 4°C on a nutator. The next day beads and supernatants were separated by centrifugation (2 min, 100g at 4°C). Beads were washed 3 times in 1 ml Triton X-100 buffer plus inhibitors at 4°C. Beads were resuspended in 2x Laemmli buffer and boiled for 5 min to elute proteins.

### RNA and RT-qPCR

Yeast RNA was prepared using the RNeasy Plus Mini Kit (Quiagen) according to the manufacturers protocol. cDNA synthesis was performed using Super Sript II Reverse Transcriptase Kit (Invitrogen). 1.5 µg of RNA was transcribed into cDNA according to the manufacturers protocol. The synthesized cDNA was diluted 1:15 in nuclease-free water. qPCR was performed with a Biorad CFX Connect RT-PCR machine using the Biorad iTaq Universal SYBR Green SuperMix and 4 µl of the cDNA 1:15 dilution were use in a 20µl reaction as a template. Primers were used at a final concentration of 0.2 µM. Sequences of primers are: *MET25F:* GCCACCACTTCTTATGTTTTCG, *MET25R:* AGCAGCAGCACCACCTTC, *GSH1F*: TGACAGCATCCATCAGGACCAG, *GSH1R:* GGAAGCCAGTTTCGCCTCTTTG, 18SrRNAF: GTGGTGCTAGCATTTGCTGGTTAT, 18SrRNAR: CGCTTACTAGGAATTCCTCGTTGAA. 18S rRNA was used as a normalization control for each sample. The ΔΔCt method was used for analysis. Data are represented as mean ±SD.

---

**Table S1.**

| Strain | relevant genotype | Source |
| --- | --- | --- |
| 15Daub | a bar1Δ ura3Δns, ade1 his2 leu2-3112 trp1-1 | Reed et al., 1985 |
| PY236 | pep4::URA3 | Kaiser et al., 2000 |
| PY1073 | 12mycMET30::ZEO pep4::URA3 | Yen et al., 2012 |
| PY1630 | CDC48RGS6xHis::KAN 12MycMET30::ZEO pep4::URA | Yen et al., 2012 |
| PY1729 | CDC48RGS6xHis::KAN 12MycMET30::ZEO Shp1::HYG pep4::URA | Yen et al., 2012 |
| PY2230 | 2xFLAGOsTir::URA Shp13xHA_AID::TRP YCpLEUprmet30_9xMYCMET30 | This Study |
| PY2231 | CDC48RGS6xHis::KAN 12MycMET30::ZEO Shp13xHA::TRP pep4::URA | This Study |
| PY2232 | CDC48RGS6xHis::KAN 12MycMET30::ZEO Shp13xHA ΔUBA pep4::URA | This Study |
| PY2234 | CDC48RGS6xHis::KAN 12MycMET30::ZEO Shp13xHA ΔCIM1::TRP pep4::URA | This Study |
| PY2235 | CDC48RGS6xHis::KAN 12MycMET30::ZEO Shp13xHA ΔCIM2::TRP pep4::URA | This Study |
| PY2236 | CDC48RGS6xHis::KAN 12MycMET30::ZEO Shp13xHA ΔUBX::TRP pep4::URA | This Study |

**Table S2.**
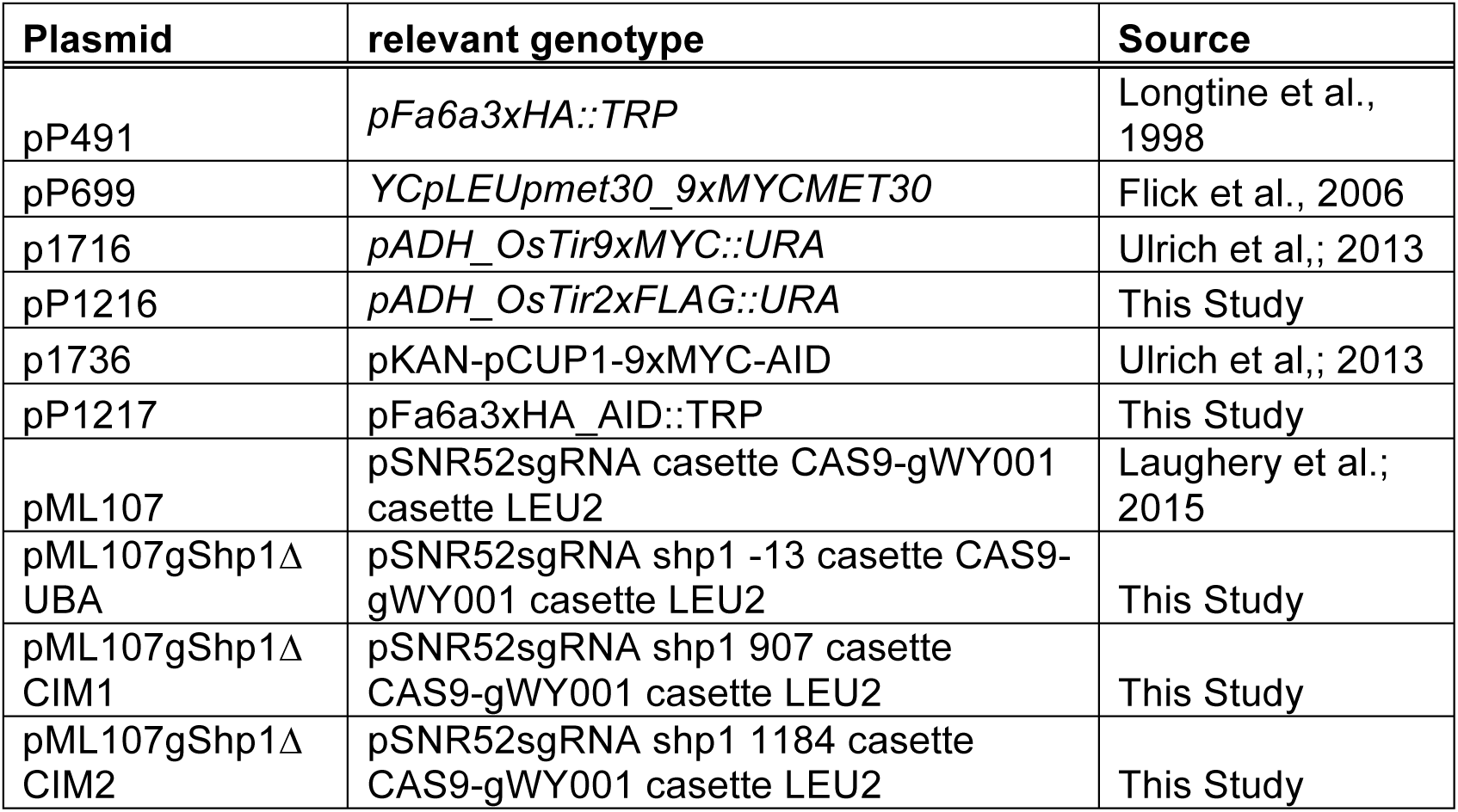

**Table S3.**
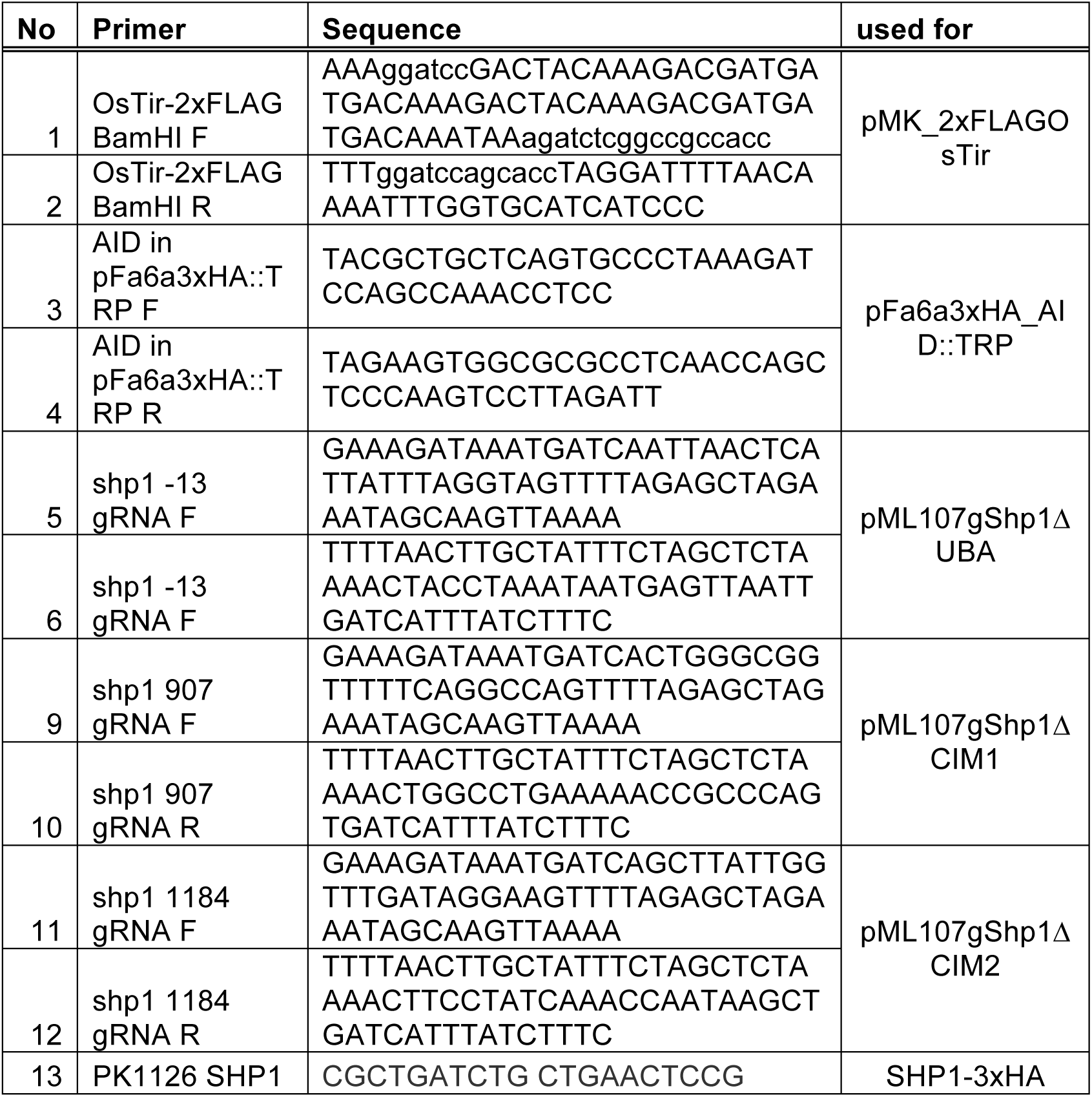
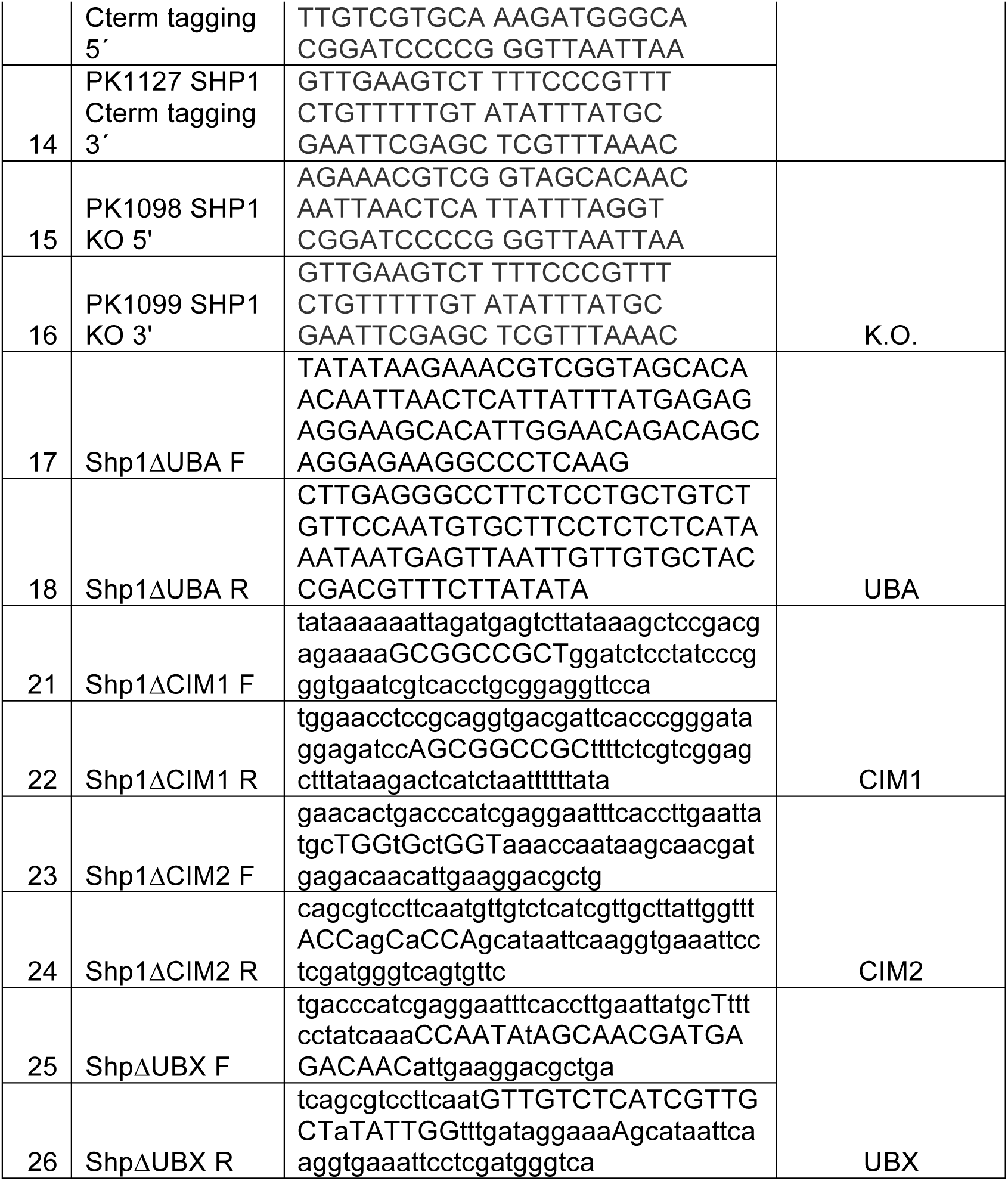

## Notes

#### Summary of Updates

Results and Discussion were updated for clarification. Figure labels were updated. Supplemental files were added.

